# Molecular basis of ubiquitination catalyzed by the bacterial transglutaminase MavC

**DOI:** 10.1101/2020.03.10.984922

**Authors:** Hongxin Guan, Jiaqi Fu, Ting Yu, Zhao-Xi Wang, Ninghai Gan, Yini Huang, Vanja Perčulija, Yu Li, Zhao-Qing Luo, Songying Ouyang

## Abstract

The *Legionella pneumophila* effector MavC is a transglutaminase that carries out atypical ubiquitination of the ubiquitin (Ub) E2 conjugation enzyme UBE2N by catalyzing the formation of an isopeptide bond between Gln40 of Ub and Lys92 (or to a less extent, Lys94) of UBE2N, which results in inhibition of UBE2N signaling in the NF-κB pathway. In the absence of UBE2N, MavC deamidates Ub at Gln40 or catalyzes self-ubiquitination. However, the mechanisms underlying these enzymatic activities of MavC are not fully understood at molecular level. In this study, we obtained the structure of the MavC-UBE2N-Ub ternary complex that represents a snapshot of covalent cross-linking of UBE2N and Ub catalyzed by MavC. The structure reveals the unique way by which the cross-linked catalytic product UBE2N-Ub binds mainly to the Insertion and the Tail domains of MavC prior to its release. Based on our structural, biochemical and mutational analyses, we proposed the catalytic mechanism for both the deamidase and the transglutaminase activities of MavC. Finally, by comparing the structures of MavC and MvcA, the homologous protein that reverses MavC-induced UBE2N ubiquitination, we identified several key regions of the two proteins responsible for their opposite enzymatic activity. Our results provide insights into the mechanisms for substrate recognition and ubiquitination mediated by MavC as well as explanations for the opposite activity of MavC and MvcA.

## Introduction

Signal transduction in cells is often mediated by posttranslational modifications (PTMs), which impact the activity of existing proteins to allow rapid responses to upstream cues. Among more than 200 different types of PTMs identified so far ^1, 2^, ubiquitination is one of the most widely used. Canonical ubiquitination requires the activities of the E1, E2 and E3 enzymes that respectively activate, conjugate and ligate the 76-residue ubiquitin (Ub) to modify proteins ^3^. Ubiquitination itself is further regulated by ubiquitination and other types of PTMs such as phosphorylation, acetylation and ADP-ribosylation that target Ub, components of the ubiquitination machinery, or both ^4^. This complex crosstalk among various PTMs allows cells to achieve better fine-tuning of their response to various stimuli, particularly under disease conditions ^4, 5^.

Pathogens have evolved diverse mechanisms to co-opt host functions to promote their fitness. One such mechanism is the acquisition of virulence factors capable of effective modulation of cellular processes by various PTMs ^6^. *Legionella pneumophila*, the causative agent of Legionnaires’ disease, is one such example. The intracellular life cycle of this bacterium utilizes the Dot/Icm type IV secretion system that injects hundreds of virulence factors known as effectors into host cells ^7, 8^. These effectors extensively modulate cell signaling hubs such as small GTPases and the Ub network to create a niche permissive for intracellular replication of the *L. pneumophila* ^9^.

Co-option of the host Ub network by *L. pneumophila* appears to be of particular importance for modulating host cellular immune process to facilitate its intracellular replication. More than 10 effectors with E3 Ub ligase activity have been identified. Although their target proteins remain elusive in most cases, theseeffectors cooperate with E1 and E2 enzymes in host cells to form active ubiquitination machineries ^10^ (**Fig. S1A**). A paradigm shift discovery was made by the study of the SidE effector family (SidEs) that includes effectors such as SdeA, which catalyze a NAD^+^-dependent ubiquitination. This mechanism involves Ub activation via ADP-ribosylation and phosphodiesterase (PDE)-mediated ligation of phosphoribosylated ubiquitin (PR-Ub) onto serine residues of substrate proteins^11–13^ (**Fig. S1B**). Interestingly, two research groups recently reported that DupA and DupB, the two highly homologous PDE domain-containing deubiquitinases from *L. pneumophila*, similarly reverse phosphoribosyl serine ubiquitination on their substrates ^14^ (**Fig. S1B**). Moreover, the activity of SidEs is regulated by SidJ, another effector which inhibits the mono-ADP-ribosyltransferase activity by calmodulin-dependent glutamylation ^15–17^ (**Fig. S1B**).

The modification of the E2 enzyme UBE2N by MavC represents another atypical ubiquitination mechanism. In this reaction, UBE2N, which exists in cell primarily as a UBE2N∼Ub conjugate linked by a thioester bond ^18, 19^, is ligated to Ub via an isopeptide bond formed between Gln40 of Ub and Lys92 (i.e. γ-glutamyl-ε-Lys bond between Ub_Gln40_ and UBE2N_Lys92_) or, to a lesser extent, Lys94 of UBE2N ^20^. This ligation is mediated by transglutamination, a reaction that does not require exogenous energy ^21^ (**Fig. S1C-D**). Analogously to other transglutaminases that function as deamidases in the absence of their target substrates ^22^, MavC uses catalytic Cys74 that is crucial for both enzymatic activities ^23^ (**Fig. S1E**). Ubiquitination at Lys92 abolishes the activity of UBE2N, which in turn curbs the formation of K63-type polyubiquitin chains through canonical ubiquitination otherwise mediated by UBE2N, E1 and UVE1, thereby inhibiting NF-κB activation ^20^ (**Fig. S1A**).

MavC and its homolog MvcA are structurally similar ^23^ to the canonical ubiquitin deamidase cycle inhibiting factor (Cif) effectors from enteropathogenic *Escherichia coli* and its homolog in *Burkholderia pseudomallei* (CHBP) ^24, 25^ (**Fig. 1**). Both MavC and MvcA have ubiquitin deamidase activity but only MavC is able to induce monoubiquitination of UBE2N. Furthermore, we recently found that MvcA counteracts the trangslutamination activity of MavC by removing ubiquitin from UBE2N-Ub ^21^. However, although the structures of MavC and its homolog MvcA have been solved ^23^ (**Fig. 1C**), the mechanism underlying transglutaminase-induced UBE2N ubiquitination by MavC and the molecular basis for their opposite catalytic activities both remain elusive. Here, by solving the structure of the MavC-UBE2N-Ub ternary complex and comparing it to other available structures of MavC and MvcA, we illustrate the structural basis for substrate recognition by MavC and the mechanism that mediates the formation of the isopeptide bond between Lys92 in UBE2N and Gln40 in Ub. In addition, structural comparison of the MavC and MvcA in their apo form and in ternary complex has allowed us to gain insights into the basis of the opposite biochemical activity exhibited by these two highly similar proteins in terms of regulation of UBE2N ubiquitination.

**Fig. 1.**
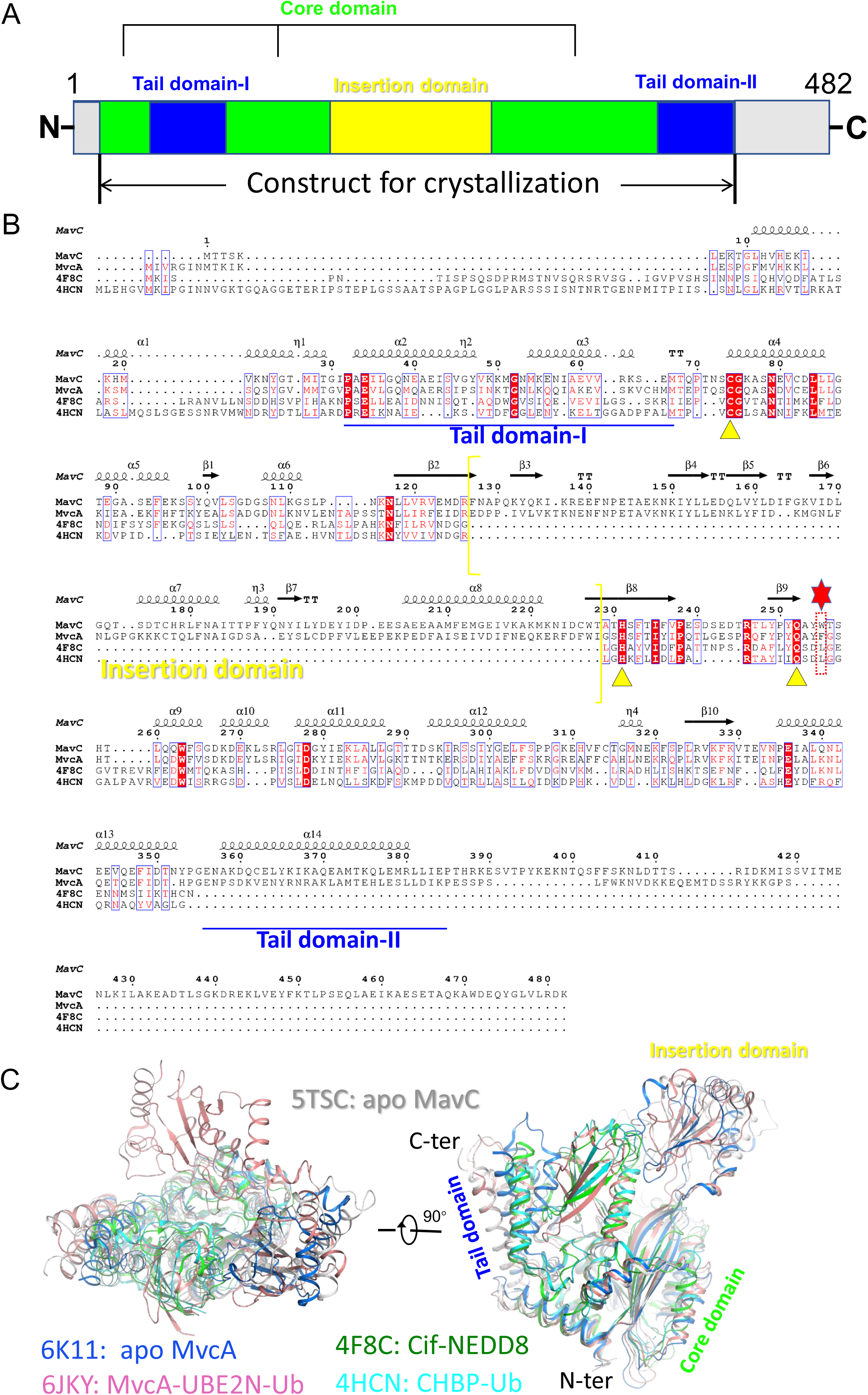
Primary sequence and three-dimensional structure comparison of MavC and its homologs MvcA, Cif and CHBP. **A.** Domain organization of MavC. The Core domain, Insertion domain and Tail domain (divided into Tail domain-I and -II) are colored green, yellow and blue, respectively. **B.** Primary sequence alignment of MavC with MvcA, Cif and CHBP generated by ClusterW (https://www.genome.jp/tools-bin/clustalw) and ESpript 3 (http://espript.ibcp.fr/ESPript/ESPript/). Every tenth residue is indicated with a dot (.) above it. Strictly conserved residues are indicated in white on a red background. The yellow triangles indicate the residues of catalytic triad sites. Residues Trp255 of MavC and Phe268 of MvcA proximal to the active site are marked by a red dotted rectangle box and a red hexagon above them. The part of sequence corresponding to the Insertion domain is enclosed by yellow brackets, whereas the parts of sequence corresponding to the Tail domain (-I and -II) are underlined by a blue line. **C.** Three-dimensional structure comparison of apo MavC (PDB ID:5TSC) with MvcA (PDB ID:6K11 and 6JKY), Cif (PDB ID: 4F8C) and CHBP (PDB ID: 4HCN) in two different orientations.

## Results

### The Insertion domain of MavC is essential for UBE2N ubiquitination but not for self-ubiquitination and ubiquitin deamidation activities of MavC

Purified MavC from *E. coli* exists primarily as a mixture of monomers and dimers in solution, both of which interact with UBE2N in size-exclusion chromatography (SEC) (**Fig. S2**). Unlike Cif and CHBP, both apo MavC (PDB ID: 5TSC) and MvcA (PDB ID: 6K11) possess a unique Insertion domain, which likely plays an important role in interaction with UBE2N ^23^ (**Fig. 1**). To test this hypothesis, we constructed a MavC truncation mutant missing Insertion domain (MavC_Δmid_, lacks residues Gln131 to Asn223) and then examined its activities (**Fig. 1A**). In contrast to wild-type MavC (MavC_WT_) that robustly induced UBE2N ubiquitination, both MavC_C74A_ and MavC_Δmid_ lost the ability to carry out UBE2N ubiquitination (**Fig. 2A left panel**). In line with the biochemical results, although MavC_Δmid_ displayed high expression levels in the *L. pneumophila* strain *ΔmavC* and was delivered into host cells, it failed to ubiquitinate UBE2N (**Fig. 2B**). Intriguingly, MavC_Δmid_ is still capable of catalyzing self-ubiquitination (**Fig. 2A right panel)** and ubiquitin deamidation (**Fig. 2C**). These results suggest that the Insertion domain is essential for UBE2N ubiquitination but not for ubiquitin deamidation and self-ubiquitination activities of MavC. Considering our findings, we further hypothesized that the Insertion domain of MavC is involved in substrate recognition.

**Fig. 2.**
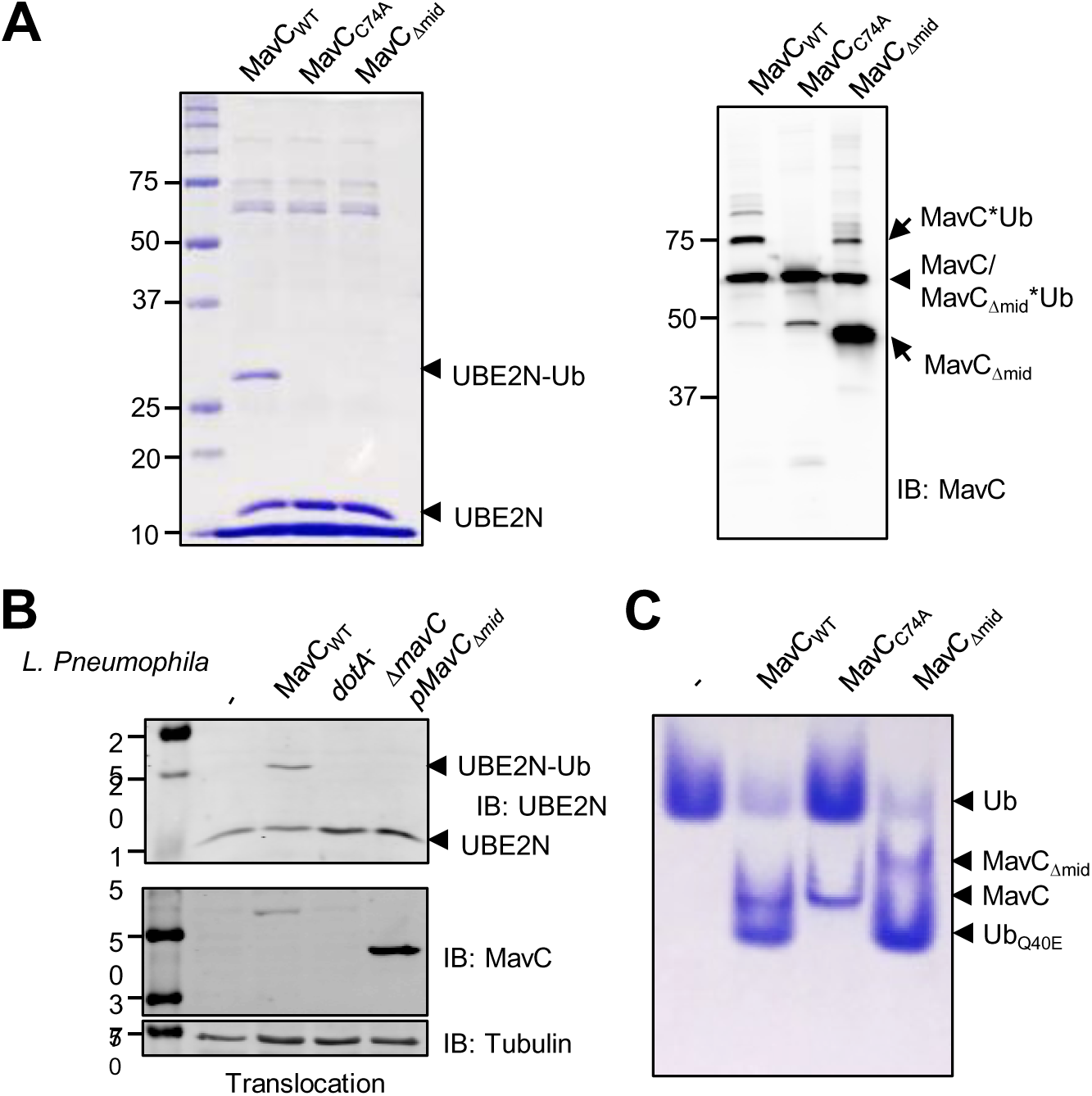
The Insertion domain of MavC is essential for UBE2N ubiquitination but not for ubiquitin deamidation and self-ubiquitination. **A.** Deletion of the Insertion domain of MavC (MavC _mid_) abolished UBE2N ubiquitination but still had no effect on ubiquitin deamidation and self-ubiquitination. Ubiquitination reactions containing MavC, MavC_C74A_ or MavC_Δmid_ were resolved by SDS-PAGE and visualized by Coomassie staining (left panel) or immunoblotting with MavC-specific antibodies (right panel). **B.** The MavC_Δmid_ mutant did not catalyze UBE2N ubiquitination in cells infected by *L. pneumophila*. U937 cells were infected with the indicated *L. pneumophila* strains for 1 h at a MOI of 10, after which the cell lysates separated by SDS-PAGE were probed by immunoblotting with the indicated antibodies. Note that the MavC_Δmid_ mutant was translocated into host cells at levels higher than wild-type bacteria. **C.** MavC mutant lacking the Insertion domain is capable of ubiquitin deamidation. Proteins in reactions containing ubiquitin and MavC or its mutants were separated by native PAGE and visualized by Coomassie staining.

### Binding affinities between MavC and its substrates

We first, examined the binding affinity between Ub and MavC. Although Ub participates in MavC-induced transglutamination *in vitro*, we could not observe the formation of a stable complex between Ub and MavC_C74A_ in solution using SEC and no detectable binding was detected in MST assays (**Fig. S3A-B**). High concentrations of Ub (>3 mM) were insufficient to generate saturation curve in MST assay. We therefore assumed that binding between these two proteins is too weak to be detected under the assay conditions. Similarly, we did not observe crystal growth when mixing MavC and Ub at 1:3 ratio. These findings are inconsistent with those of a previous study that confirmed binding between MavC and Ub by monitoring chemical shift perturbations (CSPs) in NMR titration experiments ^23^. The differences are probably due to the fact that the catalytically active MavC_WT_ used in this previous study deamidated the ^15^N-labeled ubiquitin during titration in solution nuclear magnetic resonance (NMR) ^23^. Thus, although Ub participates in MavC-mediated transglutamination and deamidation reactions, its interactions varies when assayed with enzymatically active or inactive MavC^21^.

We also examined binding between MavC_C74A_ and UBE2N with the MST assay and found that these two molecules bind at a K_d_ value of 1.14 µM in solution (**Fig. S3C**). Moreover, since the best resolution we could obtain for the MavC_C74A_-UBE2N binary complex was only about 3.5 Å, we speculated that the intermolecular interactions mediated by the Insertion domain of MavC are dynamic. Interestingly, although both MavC and MvcA contain an Insertion domain, MvcA has been showed to have no interaction with UBE2N ^23^. Taken together, these results indicate that, in spite of ∼50% sequence identity, MavC and MvcA differ in recognizing UBE2N.

To investigate the molecular basis for substrate binding and transglutaminase activity of MavC, we aimed to solve the structure of the MavC-UBE2N-Ub ternary complex. We attempted to improve the binding stability between MavC_C74A_ and its substrates Ub and UBE2N by directly supplying the product of MavC-catalyzed transglutamination on the premise that the binding affinity between MavC and cross-linked UBE2N-Ub binary complex (K_d_=1.4 µM) is similar to the binding affinity between MavC and UBE2N and would thus solve the problem of weak interactions between and MavC and free Ub (**Fig. S3D**). The MavC_C74A_ and UBE2N_K94A_ mutants were used for crystallization experiments to ensure that UBE2N is ubiquitinated only at Lys92 ^21^.

### Overall structure of MavC in complex with its product UBE2N-Ub and catalytic site interactions

By following the protocol described above, we successfully crystallized and solved the structure of the MavC_C74A_-UBE2N_K94A_-Ub ternary complex at a 2.85 Å resolution (**Table 1**). Only one copy of the ternary complex could be found in the asymmetric unit (ASU). Interestingly, the architecture of the MavC_C74A_-UBE2N_K94A_-Ub complex is highly similar to that of the MvcA-UBE2N-Ub complex reported in our previous study ^21^. The ternary complex solved in this study represents the stage of MavC-mediated UBE2N ubiquitination in which UBE2N and Ub have already been cross-linked by an isopeptide bond but are still bound to MavC (**Fig. 3A-B, Movie S1**). MavC, which assumes a concave shape, consists of a Core domain flanked by a helical Tail domain and an Insertion domain (**Fig. 1 and 3A-B**). Loops 1, 2 and 5 along with loops 3 and 4 serve as flexible hinges that respectively link the Tail domain and the Insertion domain to the Core domain, thus conferring the flexibility to the two subdomains required for binding UBE2N-Ub (**Fig. 3A-B**). The covalently linked UBE2N-Ub sits in the groove formed by the Tail domain and the Insertion domain of MavC with the UBE2N and Ub portion of the molecule hanging on either side of the enzyme.

**Fig. 3.**
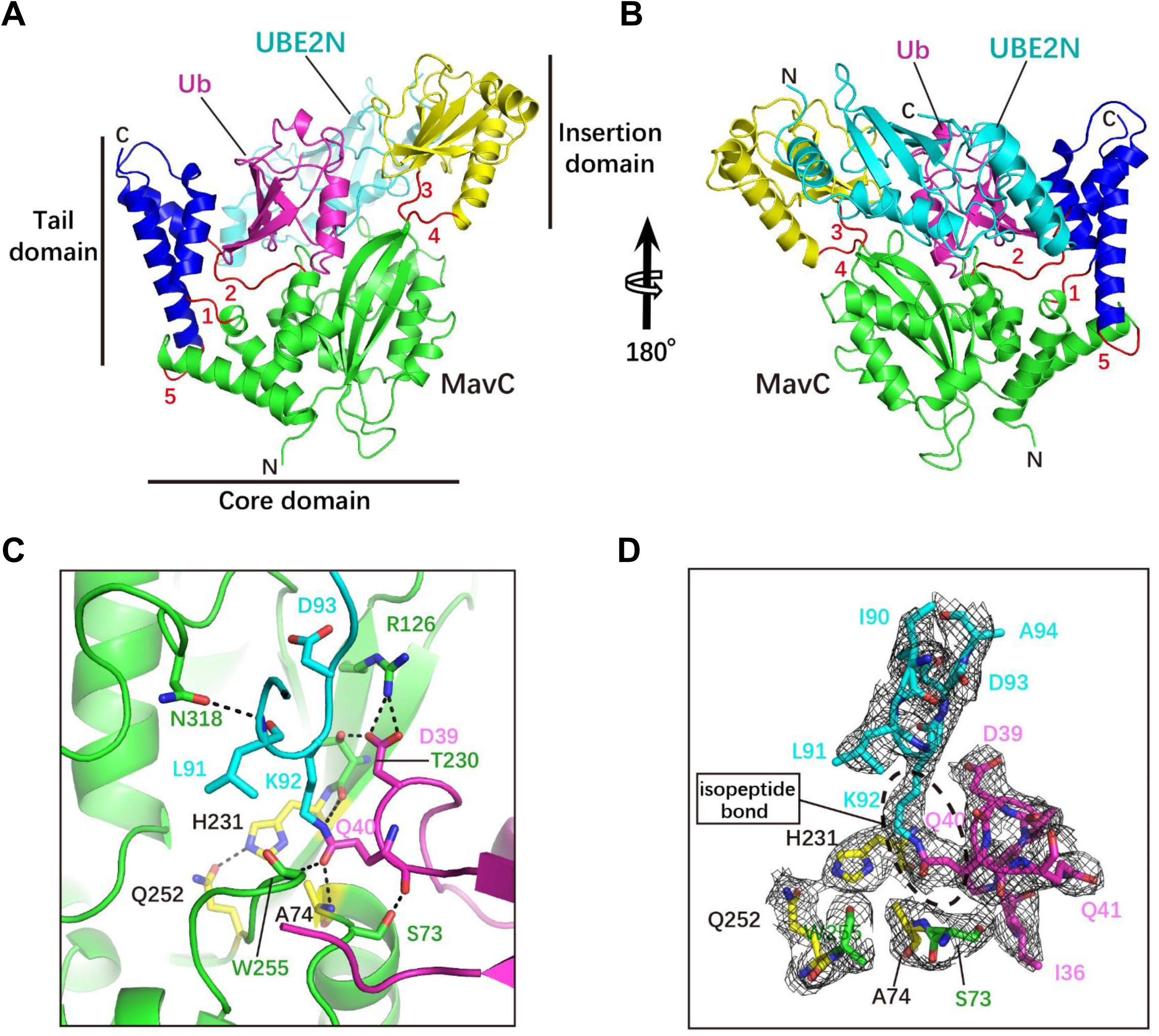
The overall structure of the MavC-UBE2N-Ub complex. **A-B**. Ribbon representation of the MavC-UBE2N-Ub ternary complex. In MavC, the Tail domain (helices α2, α3 and α14, blue) is linked to the Core domain (green) by three loops (loop 1, 2 and 5, red), whereas the Insertion domain (yellow) is linked to the Core domain by two loops (loop 3 and 4, red). Ub and UBE2N are shown in magenta and cyan, respectively. The view in panel B is generated by rotating the image in panel A by 180° around the indicated axis. **C.** MavC-induced linkage between Lys92 of UBE2N (cyan) and Gln40 of Ub (magenta). The catalytic triad (yellow) of MavC (Cys74 was mutated to Ala in our structure) and other residues participating in the reaction are shown as sticks. Hydrogen bonds are shown as dashed lines. **D.** The 2Fo-Fc map of (C) contoured at 1.2 σ. The catalytic triad (Ala(Cys)74-His231-Gln252, yellow) and Ser73 of MavC, Ile90, Leu91, Lys92, Asp93, and Ala94 of UBE2N, and Ile36, Asp39, Gln40 and Gln41 of Ub are shown as sticks.

The catalytic site for the transglutaminase is situated at the bottom of the concavity of MavC, with the Cys74-His231-Gln252 catalytic triad (Cys74 was mutated to Ala in our structure) at the center of the concave line. Several residues adjacent to the catalytic triad contribute to the stabilization of the UBE2N loop and the Ub loop that respectively contain Lys92 and Gln40, the two reactive residues covalently bonded by transglutamination. Ala74, Thr230 and Trp255 of MavC interact with Gln40 of Ub through hydrogen bonding that involves both side chain and backbone atoms. Moreover, Arg126 and Thr230 of MavC form three pairs of hydrogen bonds with Asp39 of Ub, thereby further stabilizing the Gln40-containing loop. The UBE2N loop containing Lys92 is held in place by hydrogen bonding between Asn318 of MavC and Leu91 of UBE2N (**Fig. 3D**).

### Key residues mediating interactions between MavC and UBE2N and their implications in transglutaminase of MavC

In the ternary complex, interactions between MavC and UBE2N include hydrogen bonding, electrostatic and hydrophobic interactions. Three pairs of hydrogen bonds (Lys132-Glu61, Glu207-Lys6 and Tyr198-Gln100) at the interface between the Insertion domain of MavC and UBE2N contribute to UBE2N recognition (**Fig. 4A**). Electrostatic interactions further stabilize the binding of UBE2N-Ub with MavC. These electrostatic interactions involve the loop between β6 strand and α7 helix composed of seven negatively charged residues (Asp196, Glu197, Asp200, Glu202, Glu203, Glu206 and Glu207) and three Tyr residues (Tyr189, Tyr192 and Tyr198) that interacts with a positively charged region formed by four residues (Arg6, Arg7, Lys10 and Arg14) of the α1 helix in UBE2N (**Fig. 4A**). According to the sequence conservation analysis, the above residues of MavC are conserved except Y189, E197 and Glu202 (**Fig. S4**). Hydrophobic interactions are mediated by Met317 from the Core domain of MavC which inserts into a hydrophobic pocket in UBE2N formed by residues Ile86, Leu88, Ile90, Leu99, Val104, Ile108 and Leu111 (**Fig. 4B-C**).

**Fig. 4.**
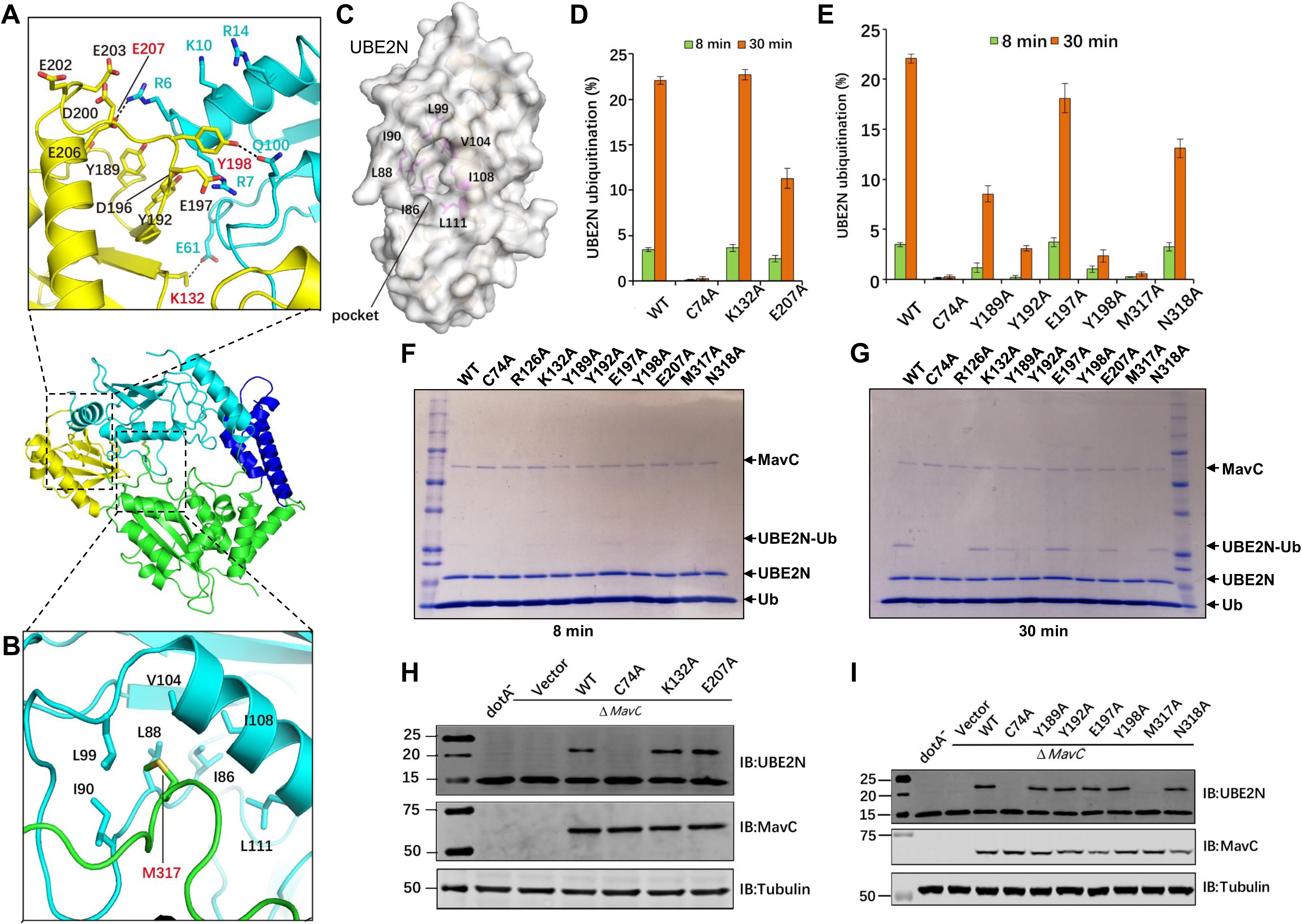
The effects of mutations in MavC on MavC-induced ubiquitination. **A-B**. Detailed views of the interactions between MavC and UBE2N in the ternary complex. Panel A: Three pairs of hydrogen bonds formed between the Insertion domain of MavC and UBE2N are shown as dashed lines, and the hydrogen-bonded residues are labeled with red numbers. Residues involved in electrostatic interactions between the negative surface of MavC and positive surface of UBE2N are labeled with black (MavC) and blue (UBE2N) numbers. The interacting residues are colored yellow (Insertion domain) and cyan (UBE2N). Panel B: The hydrophobic residues forming pocket 2 of UBE2N are shown as cyan sticks, whereas Met317 of MavC that inserts into the pocket is shown as a green stick and labeled by red number (B). **C.** The hydrophobic pocket of UBE2N (designated pocket 2) into which Met317 of MavC inserts in the ternary complex. The hydrophobic residues lining the pocket (magenta sticks) are labeled with numbers in black and UBE2N is shown as cartoon with partially transparent surface (gray). **D-E.** Mutational analysis of MavC residues involved in the MavC-UBE2N interaction interface and their importance for UBE2N ubiquitination. MavC or its mutant derivatives were added to reactions containing UBE2N and ubiquitin for 8 min or 30 min at 37°C. Samples resolved by SDS-PAGE and visualized by Coomassie staining were quantified using Image Studio. The ratios are from three independent experiments. Error bars indicate standard error of the mean (SEM). **F-G**. SDS-PAGE images of the ubiquitination assay shown in panels D-E. MavC and its mutant derivatives were added to reactions containing UBE2N and Ub, the reactions were allowed to proceed for 8 or 30 min at 37°C before SDS-PAGE and visualization with Coomassie staining. **H-I.** The ability of MavC mutant derivatives to catalyze UBE2N ubiquitination during *L. pneumophila* infection. Plasmids carrying *mavC* mutants were introduced into the Δ*mavC* mutant and the bacteria were used to infect U937 cells. Saponin-soluble lysates of infected cells resolved by SDS-PAGE were probed with antibodies specific for MavC or UBE2N.

To determine the impact of the residues that play role in interaction and recognition of UBE2N during ubiquitination, we designed a set of MavC mutants and tested their ability to produce UBE2N-Ub. Mutations of Tyr192, Tyr198 and Met317 to alanine severely impaired ubiquitination of UBE2N, whereas the Y189A and N318A mutants were only partially defective. Furthermore, mutation to alanine of E207 that hydrogen bonds with Lys6 of UBE2N caused a slight defect in catalyzing UBE2N ubiquitination, suggesting that this residue is required for optimal ubiquitination activity. In contrast, we did not detect defects associated with MavC_E197A_ (**Fig. 4D-E**). These observations are consistent with the binding results, which showed that MavC_Y192A_ and MavC_Y198A_ failed to bind UBE2N (**Fig. 4F-G**). When introduced into the *L. pneumophila ΔmavC* mutant on a plasmid, each of the abovementioned mutants can be expressed and translocated into host cells at levels comparable to that of wild-type MavC. Our in vivo results, reveal that only the M317A mutant has lost the ability to induce UBE2N ubiquitination, thus implying an important role of this residue in the transglutaminase activity of MavC (**Fig. 4H-I**). Therefore, electrostatic interactions and hydrophobic interactions mediated by Met317 are the main force to stabilize the binding of UBE2N-Ub with MavC.

To further validate the substrate recognition mechanism implied by the ternary structure, we designed UBE2N_R6A/R7A_, UBE2N_K10A/R14A_ and UBE2N_ΔN6-14_ (deletion mutant missing the residues from Arg6 to Arg14) mutants predicted to affect intermolecular electrostatic interactions and tested them for ubiquitination using the method described above. We found that neither UBE2N_R6A/R7A_ nor UBE2N_ΔN6-14_ demonstrated detectable ubiquitination and that UBE2N_K10A/R14A_ can still be ubiquitinated but at a markedly reduced level (**Fig. 5A-D**).

**Fig. 5.**
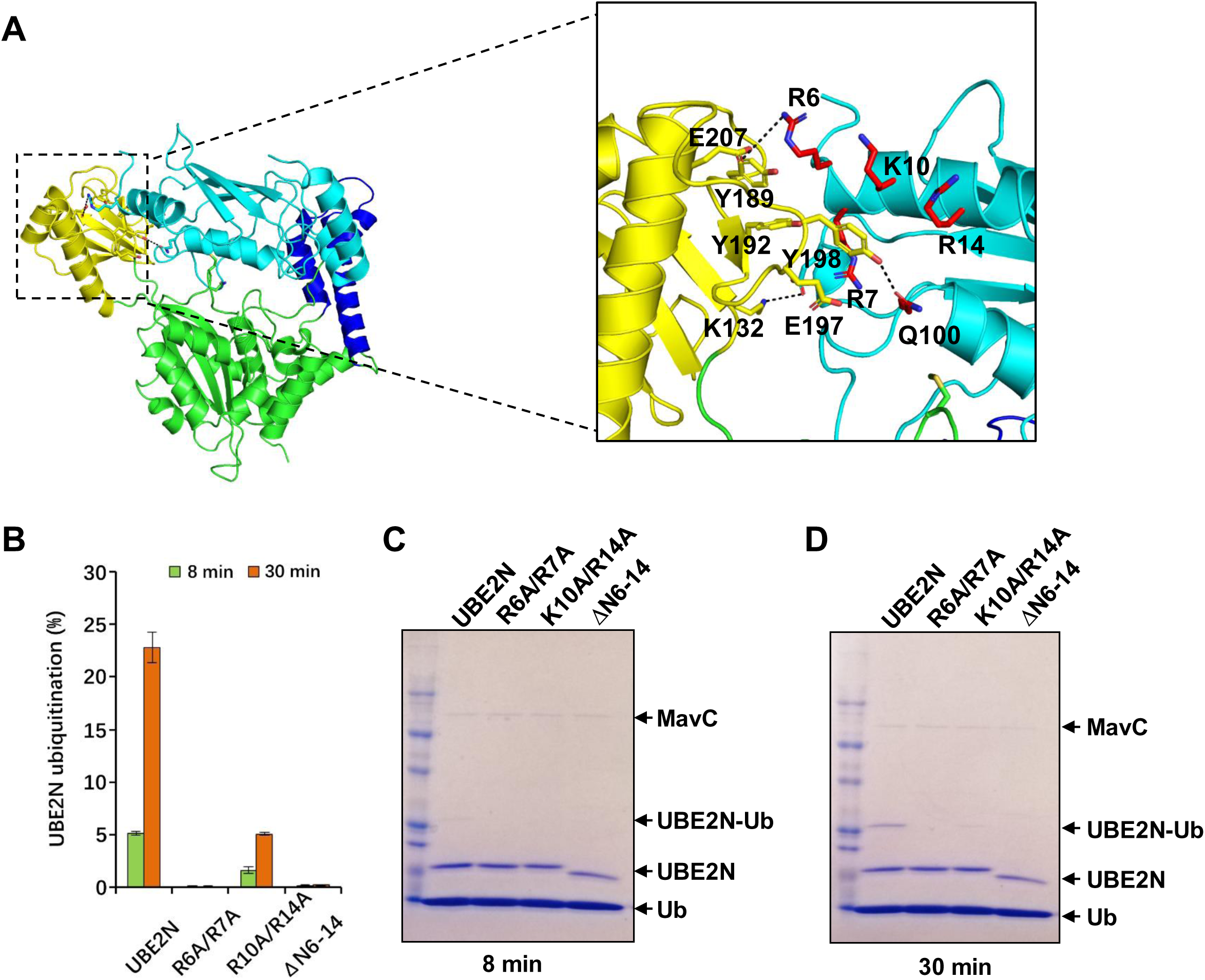
The effects of mutations in UBE2N on MavC-induced ubiquitination. **A.** Detailed views of the interactions between MavC and UBE2N in the ternary complex, similar to Fig. 4A. **B.** The effects of mutations in UBE2N on MavC-induced ubiquitination. MavC was added to reactions containing ubiquitin, UBE2N or its mutants. The reactions were allowed to proceed for for 8 min or 30 min at 37°C. Samples resolved by SDS-PAGE and visualized by Coomassie staining were quantified using Image Studio. The ratios are from three independent experiments. Error bars indicate standard error of the mean (SEM). **C-D.** The effects of mutations in UBE2N on MavC-induced ubiquitination. MavC was added to reactions containing ubiquitin, UBE2N or its mutants. The reactions were allowed to proceed for for 8 min or 30 min at 37°C before SDS-PAGE and detection by Coomassie staining.

### Interactions between MavC and Ub

Although MavC_C74A_ was not found to interact with Ub directly in SEC and does not exhibit observable affinity for it in MST assays, the structure of our ternary complex structure clearly shows that MavC_C74A_ indeed interacts with Ub via five distinct contact regions (termed contact regions A–D and the carboxyl terminus contact region CTC) on the Tail domain of MavC. Several pairs of hydrogen bonds and hydrophobic interactions from these regions contribute to positioning of the ubiquitin molecule optimal for transglutamination (**Fig. S5A-B**). Contact region A is formed by a hydrogen bond between Glu42 of MavC and Lys6 of ubiquitin (**Fig. S5C**). Contact region B involves the loop of the N-terminal β-hairpin in Ub, particularly residues Leu8, Thr9 and Gly10, which engage in hydrophobic interactions with a hydrophobic pocket of MavC formed by residues Ile31, Pro32, Ile35, Leu36 (**Fig. S5D**). In contact region C, three pairs of hydrogen bonds are formed, namely Gln31(Ub):Glu123(MavC), Glu34(Ub):Asn79(MavC) and Gly35(Ub):Arg121(MavC) (**Fig. S5E**). Contact region D features interactions mediated by a single pair of hydrogen bonds formed between Asn25 of ubiquitin and Ile163 of MavC (**Fig. S5F**). Finally, the flexible tail at the carboxyl terminus of ubiquitin is held in place by contacting an adjacent groove of MavC through hydrogen-bonding involving backbone atoms of Glu66, Gln69, His258 and side chains of Arg72 and Arg74 of the ubiquitin tail (**Fig. S5G**).

### Rotation of the Insertion and Tail domains of MavC is essential for delivering UBE2N and Ub to the trangslutaminase active site

In comparison to the apo form, the Insertion domain of MavC underwent a distinct approximately 30° anticlockwise rotation in the ternary complex. This movement allowed formation of electrostatic interactions and hydrogen bonds between the Insertion domain and positively charged region of UBE2N (**Fig. 6A-C**). The loop 1 (Leu88-Trp95) of UBE2N and the loop2 (His311-Lys320) of MavC interlock and thus strengthen binding between MavC and UBE2N and firmly stabilize the Lys92 of UBE2N in the enzyme activation center (**Fig. 6D-E, Movie S2**). Similarly to Insertion domain, the Tail domain of MavC also rotates anticlockwise relative to its position in apo form MavC. The rotation of the Tail domain instigates hydrogen bonding between Gln42 of MavC and Lys6 of Ub and electrostatic interactions (**Fig. 6F-G**). In addition, the loop 3 (Thr71-Ser73) and loop 4 (Gln252-Ser257) play an important role in maintaining the Lys92 of UBE2N and Gln40 of Ub in the precise location of the enzyme activation center (**Fig. 6H**). Based on these findings, we propose that a anticlockwise rotation of the Insertion and Tail domains in MavC during ternary complex formation moves Lys92 of UBE2N and Gln40 of Ub closer to the catalytic center, and the Core domain serves as a scaffold to stabilize UBE2N_Lys92_ and Ub_Gln40_ in the catalytic center, these three domains complete the transglutamination reaction cooperatively.

**Fig. 6.**
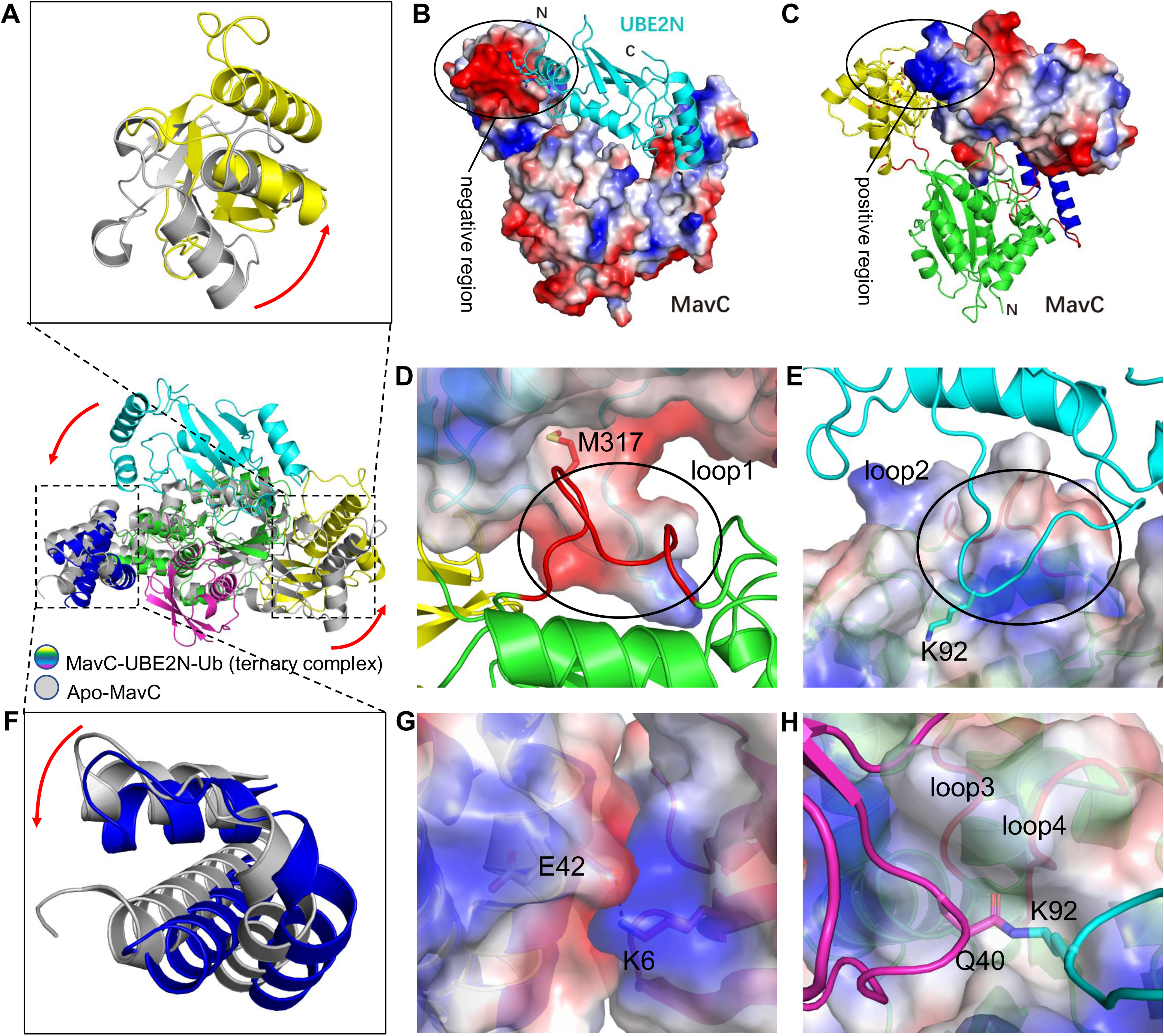
Structural details of UBE2N and Ub recognition by MavC. **A.** Superimposition of Insertion domains of UBE2N-Ub bound and apo MavC. The rotation of the Insertion domain from its position in apo MavC to its position in the ternary complex is indicated by an arrow. **B-C.** The Insertion domain of MavC in the ternary complex undergoes a rotation during the catalysis that links UBE2N and Ub. Positively charged residues of UBE2N (cyan cartoon) and the negatively charged region of MavC are indicated by a circle (B). Negatively charged residues of the Insertion domain of MavC-UBE2N-Ub (yellow cartoon) and positively charged region of UBE2N (cyan cartoon) are indicated by a circle (C). **D-E.** Two loops stabilize the binding of UBE2N to MavC. Loop1 (Leu88-Trp95, red cartoon) of MavC is indicated by a circle. Met317 of MavC is shown as red stick which inserts into the pocket of UBE2N electrostatic surface (D). Loop2 (His311-Lys320) of UBE2N is indicated by a circle. Lys92 of UBE2N is shown as cyan stick which inserts into the pocket of ternary complex electrostatic surface (E). **F.** Superimposition of Tail domains of UBE2N-Ub bound and apo MavC. The rotation of the Tail domain from its position in apo MavC to its position in the ternary complex is indicated by an arrow. **G.** The interacting interface of MavC-UBE2N-Ub Tail domain and Ub are shown as electrostatic surface. Negatively charged Glu42 and positively charged Lys6 are shown as sticks. **H.** Two loops stabilize the binding of isopetide bond between UBE2N and Ub to MavC. The isopetide bond between UBE2N (cyan) and Ub (magenta) is shown as sticks. Two hydrophobic loops (loop3 and loop4) are shown as red cartoon and labeled on the outer side of the electrostatic surface.

### The mechanism for MavC-mediated transglutamination and molecular basis of opposite enzymatic activities of MavC and MvcA

MvcA is a deubiquitinase that counteracts the activity of MavC by removing ubiquitin from UBE2N-Ub ^21^. However, in spite of their opposite enzymatic activities, these two proteins share approximately 50% identity and their structures are highly similar (**Fig. 1C**). Apo forms of MavC and MvcA represent the closed catalytically inactive form of the two proteins characterized by high conformational stability, which can be seen when superimposing three available structures of apo MvcA (PDB ID: 5SUJ ^23^, 5YM9, and also our previous work with PDB ID 6K11 ^21^) — the three structures and all monomers in an asymmetrical unit align with maximal RMSD of 0.613 Å (**Fig. 1C**). Hence, we used the apo form structures as reliable reference point in our attempt to determine functional divergence of MavC and MvcA by comparing the structures of MavC and MvcA in their apo form and in ternary complexes. Interestingly, the structural similarity between MavC and MvcA from respective ternary complexes is markedly higher than between apo-MavC and apo-MvcA (**Fig. S6A-D**). This suggests that the two proteins undergo significant conformational changes during transition to ternary complexes. In both proteins, these conformational changes can be mainly contributed to rotational movement of the Insertion domain and Tail domain relative to the Core domain.

To investigate whether the opposite enzymatic activities are the consequence of divergent catalytic reactions, we searched for potential structural differences in the catalytic triads of MavC and MvcA. The two proteins utilize an identical Cys-His-Glu catalytic triad in which the catalytic cysteine (Cys74 in MavC and Cys83 in MvcA) is essential both for the formation and the cleavage of the isopeptide bond between Lys92 of UBE2N and Gln40 of Ub ^20, 21^. In a comparable scenario, SdeA and DupA utilize an identical set of catalytic residues, but conformational differences between these residues give rise to opposite activities in phosphoribyl ubiquitination ^26^. As inferred by comparison of the apo-MavC and apo-MvcA structures, the catalytic triads (Cys, His, Glu) of MavC and MvcA assume equivalent conformations in the inactive state (**Fig. 7A**). Interestingly, the comparison of MavC-UBE2N-Ub and MvcA-UBE2N-Ub ternary complexes showed that all three residues in the catalytic triads are in similarly close alignment, and the differences in positioning of the isopeptide bond between Lys92 of UBE2N and Gln40 of Ub are minute (**Fig. 7B**). We therefore reasoned that transglutamination and deubiquitination mediated by MavC/MvcA are in fact forward and backward reactions of the same reversible catalytic reaction, respectively and that the preference for the direction in which the reaction proceeds is governed by other factors such as substrate binding affinity and stability of the substrate/product bound state. These observations and the fact that the Insertion domain and Tail domain undergo rotational movement during formation of the MavC/MvcA ternary complex prompted us to divert our attention to these two domains in further analysis.

**Fig. 7.**
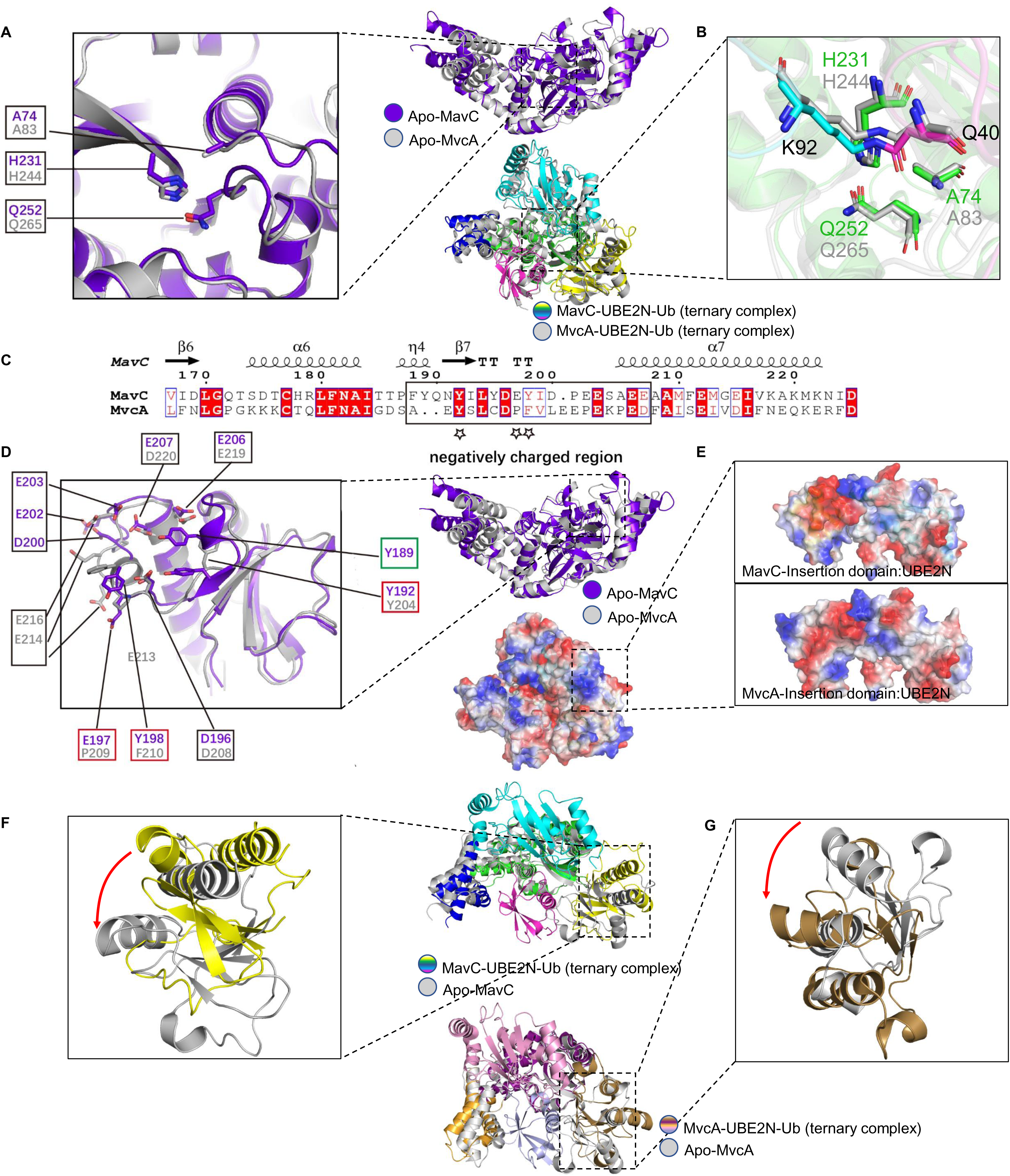
Structural comparison and analysis of Insertion domains and catalytic triads of MavC and MvcA. **A-B**. Superimposition of the catalytic triads (panel A) of Apo-MavC (purple) and Apo-MvcA (gray). Superimposition of the catalytic triads (panel B) of MavC-UBE2N-Ub (colorized) and MavC-UBE2N-Ub (gray) ternary complexes. The catalytic triads of MavC and MvcA are shown as sticks. **C.** Structure-based sequence alignment of the loop between β6 and α7 that contains negatively charged amino acids. The negatively charged region is boxed by black rectangle, and the key residues involved in the interactions between MavC and UBE2N are labeled by stars. **D-E.** Structural comparison of Insertion domains of MavC and MvcA. The residues involved in the formation of the negatively charged region are shown as sticks, corresponding residues in MavC and MvcA are labeled by black boxes, and the key residues are highlighted by red boxes (panel D). Superimposition of the Insertion domains of MavC-UBE2N-Ub and MavC-UBE2N-Ub ternary complexes. The Insertion domains of MavC and MvcA are shown as electrostatic surface (panel E). **F-G.** Superimposition of Insertion domains of MavC-UBE2N-Ub ternary complex (colorized) and apo MavC (gray) (panel F), and MvcA -UBE2N-Ub ternary complex (colorized) and apo MvcA (gray) (panel G). The rotation of the Insertion domains from their position in the apo protein to their position in ternary complex is indicated by an arrow.

While analyzing the Insertion domains of MavC and MvcA, we noticed that their negatively charged regions vary considerably in terms of negatively charged amino acid composition (**Fig. 7C**). Specifically, the negatively charged region of MavC Insertion domain contains seven negatively charged amino acids, whereas the corresponding region in MvcA contains only five, indicating that the net electric charge of the negatively charged region in MavC is more negative than the one in MvcA (**Fig. 7D).** In both ternary complexes, the Insertion domain utilizes the negatively charged region to interact with the positively charged region of UBE2N through electrostatic forces (**Fig. 7E**). Moreover, the conformational changes observed between the two ternary complexes and their apo form structures imply that the Insertion domains of both proteins bind UBE2N via anticlockwise rotation (**Fig. 7F-G**). Thus, the interaction forces and mechanisms for UBE2N by MavC and MvcA are highly similar, but the binding affinity between UBE2N and MvcA is likely to be lower than between UBE2N and MavC because of lower net negative charge.

As one of the substrates, Ub mainly interacts with the Tail domain of the MavC and MvcA. Intriguingly, comparison of the MavC and MvcA ternary complexes to their apo forms shows that the Tail domain of MavC rotates anticlockwise and the Tail domain of MvcA rotates clockwise (**Fig. 8A-B)**. Since the surface of Ub that interacts with the Tail domains of MavC and McvA is positively charged, the anticlockwise rotation of MavC Tail domain increases local negative potential, which likely results in tighter binding with Ub (**Fig. 8C-E**). On the contrary, the clockwise rotation of MvcA Tail domain creates an obvious positive bulge, which probably facilitates uncoupling of Ub from UBE2N (**Fig. 8F-H**). Another difference between MavC and MvcA is a residue in proximity to the active site, i.e. Trp255 of MavC and the corresponding residue Phe268 in MvcA (**Fig. 1B, 8I-J, S4**). The distinct properties of these two residues alter substrate binding in the active site as well as orientations of the Tail and Insertion domains that contact Ub and UBE2N.

**Fig. 8.**
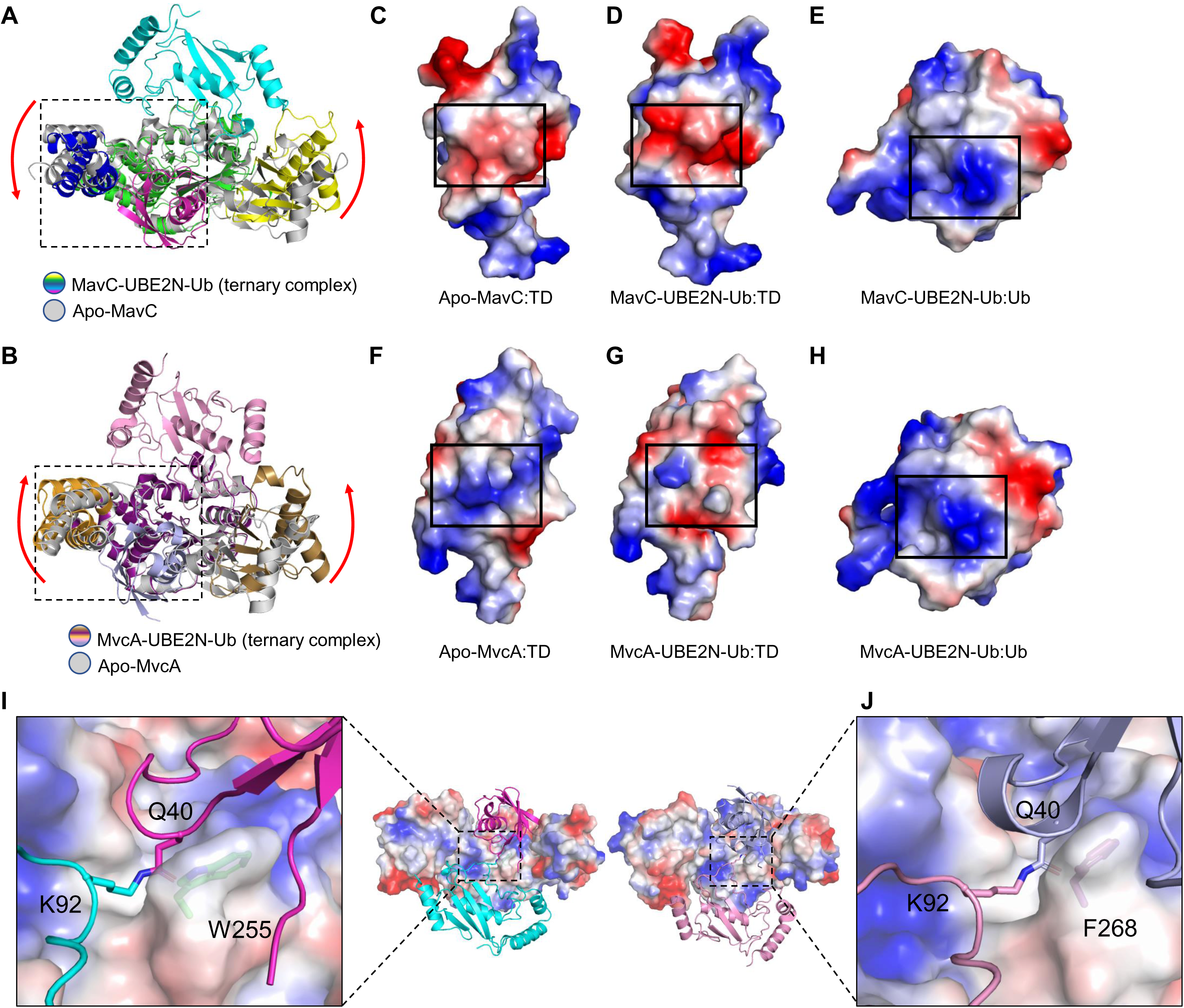
Structural comparison and analysis of Tail domains and catalytic triads of MavC and MvcA. **A.** Superimposition of MavC-UBE2N-Ub (colorized) and apo MavC (gray). The rotations of the Insertion domain and Tail domain from their positions in the apo MavC to their position in the ternary complex are indicated by red arrows. **C-E.** Electrostatic surfaces of apo MavC Tail domain, MavC-UBE2N-Ub Tail domain and Ub. **F-H.** Electrostatic surfaces of apo MvcA Tail domain, MvcA -UBE2N-Ub Tail domain and Ub. **I-J.** MavC and MvcA from their ternary complex are shown as electrostatic surfaces. The isopeptide bond (cyan and magenta) in MavC-UBE2N-Ub and isopeptide bond (light blue and pink) in MavC-UBE2N-Ub are shown as sticks. Trp255 of MavC and Phe268 of MvcA in the active site are shown as ticks and labeled on the outer side of the electrostatic surface.

In conclusion, we summarize all the structural analyses described above to propose the molecular basis for the opposite enzymatic activities of MavC and MvcA in terms of regulation of UBE2N ubiquitination (**Fig. 9A-B)**. The initial substrate engagement of MavC/MvcA involves binding of the UBE2N∼Ub conjugate (the primary form in which UBE2N exists in cell ^18, 19^)/cross-linked UBE2N-Ub through interactions that induce rotation of the Insertion domain and Tail domain, which in turn positions UBE2N∼Ub/ UBE2N-Ub in the active site of the respective protein. In MavC, the rotation of the Insertion domain and the Tail domain results in more stable binding between the UBE2N∼Ub conjugate and MavC, which is favorable for transglutamination (**Fig. 9A**). Conversely, the rotation of the Tail domain in MvcA and lower binding affinity of the Insertion domain for UBE2N promote the dissociation of Ub and UBE2N from the enzyme, thus leading to deubiquitination (**Fig. 9B**).

**Fig. 9.**
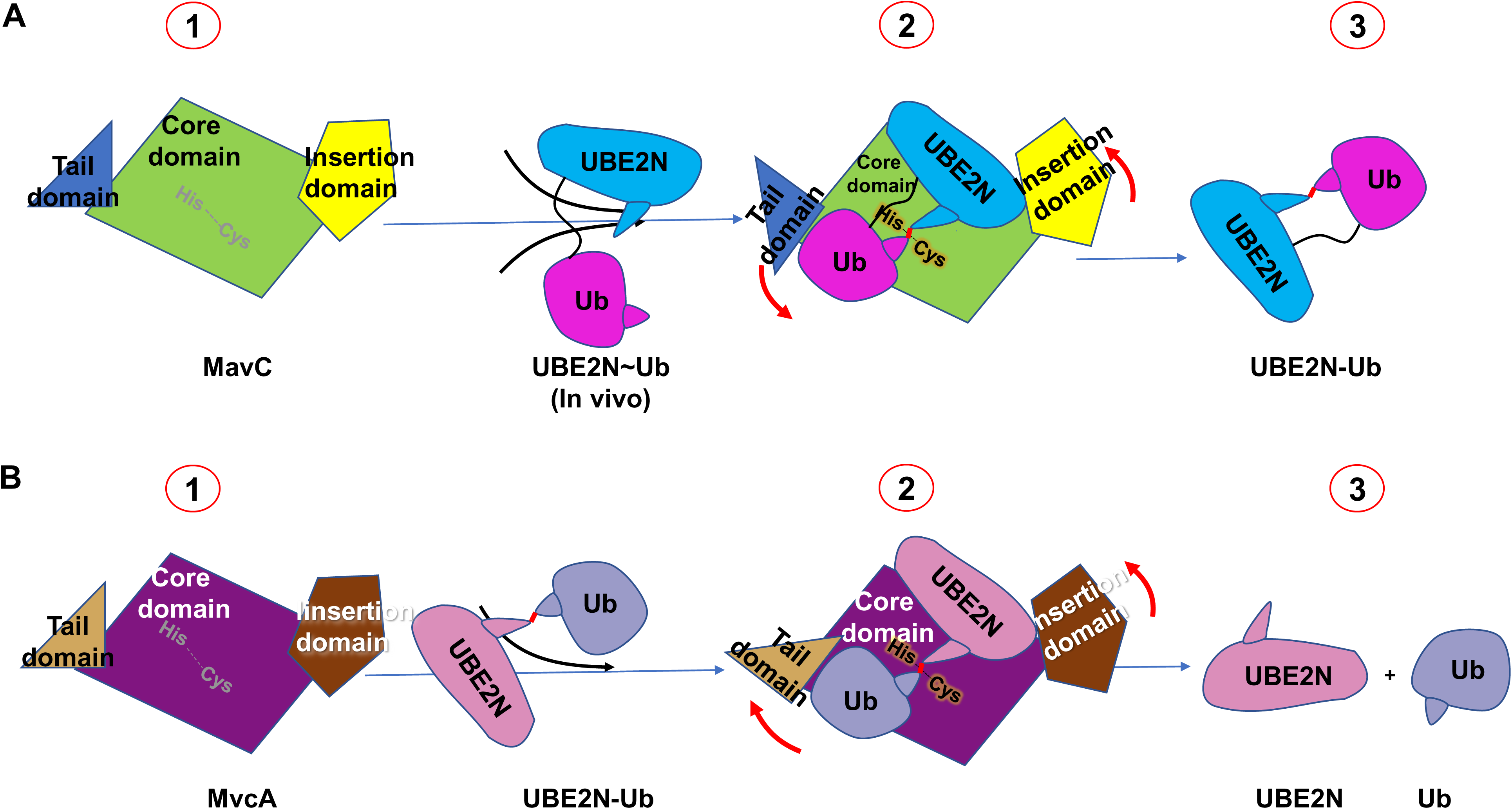
Models for mechanisms underlying MavC trangslutaminase and MvcA deubiquitination activities. **A-B.** UBE2N∼Ub binds to MavC in cells (A) and UBE2N-Ub binds to MvcA (B) by rotation of their Insertion domains and Tail domains in directions indicated by red arrows (1-2). Products are released and the catalytic reactions are completed (3).

## Discussion

MavC is a transglutaminase that catalyzes covalent cross-linking of Gln40 of ubiquitin to either Lys94 or Lys92 of UBE2N (**Fig. S1C**), and it also possess deamidase activity that targets Ub at Gln40 in the absence of UBE2N ^23^(**Fig. S1D**). The effects of MavC is counteracted by MvcA, which is a close homolog of MavC that functions as a deubiquitase against UBE2N-Ub ^21^(**Fig. S1D**). Like MavC, MvcA also exhibits ubiquitin deamidase activity in a manner similar to such bacterial deamidase such as Cif and CHBP that deamidate Gln40 of Ub and Ub/NED88, respectively ^24, 25^. Structural comparison of these four enzymes reveals that the structures of CHBP and Cif align strikingly well with the Core and Tail domains of MavC and MvcA (**Fig. 1**). However, the Insertion domain appears to be characteristic of MavC and MvcA as its structural equivalent is absent in CHBP or Cif (**Fig. 1**). Therefore, we reason that the Core and Tail domains of MavC and MvcA are sufficient for their deamination activity. Indeed, deletion of the Insertion domain abolishes the transglutamination activity of MavC while remains largely active in its deamination activity (**Fig. 2)**.

That being said, what is the functional significance of the Insertion domain? CHBP is known to target multiple signaling pathways with its ubiquitin deamidation activity via blocking free ubiquitin chain synthesis by different E3-E2 pairs, leading to the inhibition of ubiquitination of RhoA mediated by a Cullin-based E3 complex, and subsequently cell cycle arrest ^27, 28^. In contrast, the scope of activity for MvcA and MavC appears narrower as they specifically regulate ubiquitination in UBE2N-related pathways such as NF-κB signaling ^29^. Previously reported structure of the MvcA-UBE2N-Ub ternary complex reveals that the Insertion domain is involved in recognition of the UBE2N-Ub substrate ^21^. In line with that, our study shows that the Insertion domain of MavC is also involved in the recognition of UBE2N ^20^. Hence, the Insertion domain enables MvcA and MavC to act specifically on UBE2N, thereby making the regulation of host ubiquitination signaling by MavC and MvcA more specific and precise.

Catalytic domains that mediate chemically opposite reactions in highly homologous proteins are not unprecedented: the PDE domains of DupA/B and SidE enzymes are another example. PDE domains of SidE enzymes have moderate binding affinity for Ub and catalyze PR ubiquitination, whereas PDE domains of DupA/B bind strongly to Ub and mediate the removal of PR-Ub. The formation of stable enzyme-substrate complexes is required in mediate cleavage reaction, while the ligation reaction requires moderate binding affinity to substrates, probably allowing efficient release of products ^26^. MavC and MvcA are highly similar proteins with approximately 50% sequence identity and identical catalytic triad (**Fig. 1B**), yet MavC primarily catalyzes the formation of an isopeptide bond between UBE2N_Lys92_ and Ub_Gln40_, whereas MvcA is responsible for breaking of the same isopeptide bond. The structure of MavC ternary complex shows that the negatively charged surface of Tail domain attracts the positively charged surface of Ub, resulting in stable binding between MavC and Ub. Concurrently, the anticlockwise rotation of the Ub-bound Tail domain draws Ub closer to UBE2N and the active center to facilitate isopeptide bond formation. In MvcA, however, the surface charge of Tail domain is repulsive to the Ub and low binding affinity of UBE2N favors the dissociation of Ub from MvcA, which in turn facilitates the isopeptide bond breaking and separation of Ub and UBE2N from MvcA. Thus, given their overall structural similarity and equivalent chemical nature of the ubiquitin ligation by transglutamination and deubiquitination activities, the fact that MavC and MvcA are licensed with opposite enzymatic activities is due to differences in substrate binding preference and stability of the substrate/product-bound intermediates. We proposed that the formation of stable enzyme-substrate complexes is required for the synthesis reaction, whereas weak interactions between enzyme and product favor cleavage reactions.

For the transglutaminase activity of MavC, the Insertion domain enables the specific binding of MavC to UBE2N, whereas the Tail domain maintains the binding with Ub, and the Core domain serves as scaffold to stabilizes UBE2N_Lys92_ and Ub_Gln40_ in the catalytic center. These three domains coordinate to complete the catalytic reaction. The shape complementarity and architecture, rather than specific individual interactions, is crucial for isopeptide formation of UBE2N and Ub. For the deamidase activity of MavC, the Insertion domain is dispensible. In the catalyzing process of MavC, Cys74 first attacks Gln40 in ubiquitin to form a thioester intermediate. When UBE2N is present, the acylated MavC reacts with the amine donor from the ε -lysine in UBE2N to form an intermolecular isopeptide bond. In the absence of UBE2N, the acylated MavC is further attacked by a nucleophilic water to produce UbQ40E. MvcA can deubiquitinate the UBE2N-Ub product from MavC transglutaminase activity ^20, 21^(**Fig. S7**).

As a key checkpoint for activation of the NF-κB pathway, UBE2N appears to be a common target of such cellular subversion as bacterial infection; it can be regulated by diverse mechanisms such as deamidation and ISGylation ^30 4^. MavC and MvcA regulate the activity of UBE2N in spatial and temporal manner via their opposite enzymatic activities. Cross-linking of UBE2N and Ub catalyzed by MavC leads to the inhibition of the UBE2N-mediated NF-κB activation, which is essential in the early infection, whereas the deubiquitinase activity of MvcA restores the activity of UBE2N to allow the recovery of host signaling pathways in the later phases of infection when the bacteria have already successfully evaded host detection. Such mechanisms may allow less disruptions to the pathogen or promote symbiotic coexistence between the pathogen and their hosts under conditions of evolutionary pressure.

## Supporting information

Manuscript File

## Data availability

Coordinates and structure factors have been deposited in the Protein Data Bank (PDB) under accession numbers 6KL4.

## Acknowledgements

This work was supported by the National Natural Science Foundation of China grants 31770948, 31570875 (SO) and 31900879 (HG), the National Institutes of Health grant R01AI127465 (ZQL), Marine Economic Development Special Fund of Fujian Province (FJHJF-L-2020-2) and the High-level personnel introduction grant of Fujian Normal University (Z0210509). This work was also supported by the scientific research innovation program “Xiyuanjiang River Scholarship” of the College of Life Sciences, Fujian Normal University. The diffraction data were collected at the beamline BL-17U1 of Shanghai Synchrotron Radiation Facility (SSRF).

## Author contributions

SO and ZQL conceived the project. HG, TY and YH crystalized the complexes and determined the structures; YL made the movie; JF and NG constructed and analyzed the mutants; HG, ZQL, JF and SO analyzed the data. HG, JF, S.O., and ZQL wrote the manuscript.

---

**Fig. S1.**
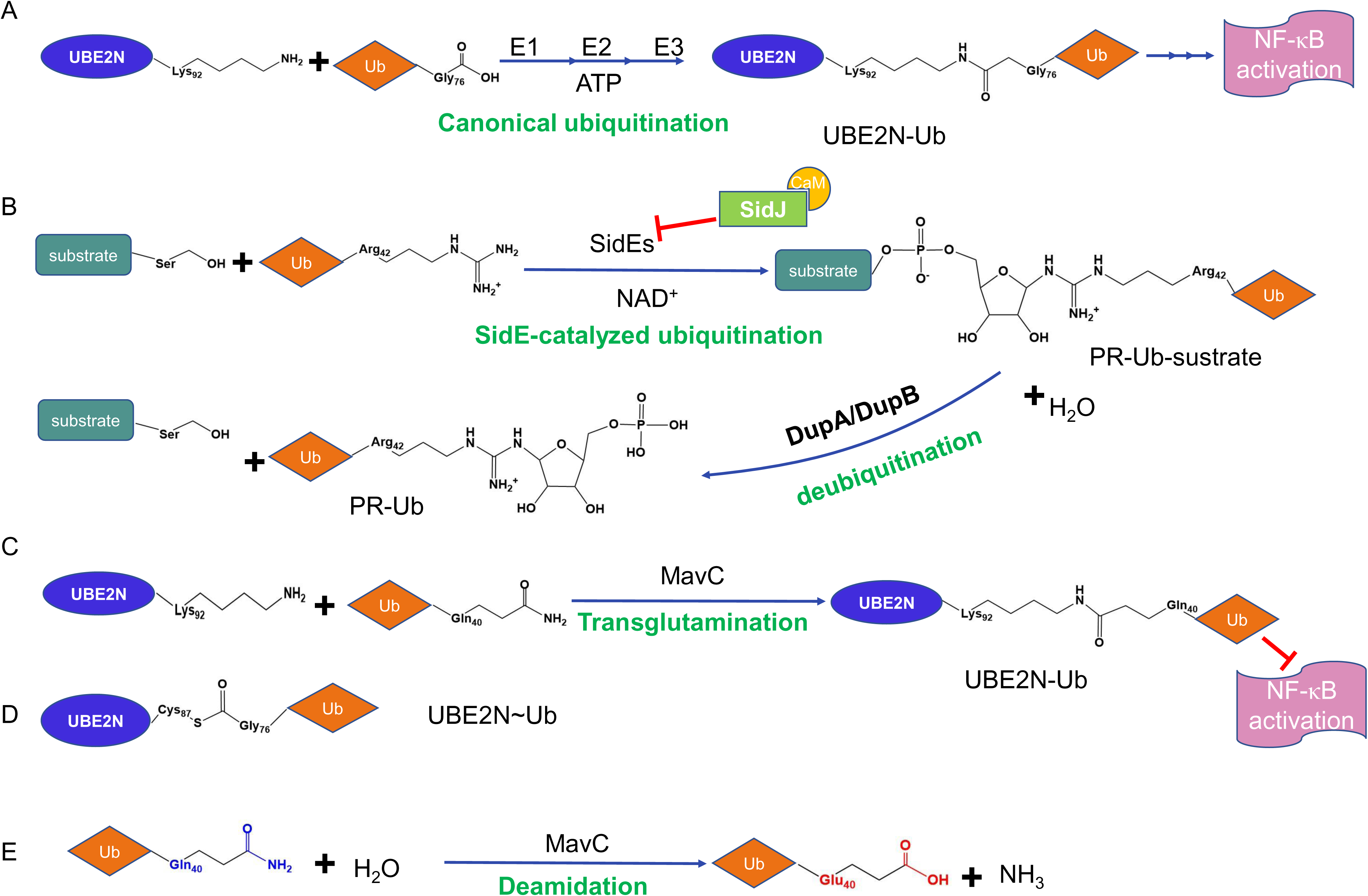
Mechanisms of ubiquitination and deubiquitination. **A.** General scheme of canonical ubiquitination. The product UBE2N-Ub suppresses the activation of NF-κB. **B.** SidE-catalyzed ubiquitination that is negatively regulated by SidJ/CaM and DupA/DupB-catalyzed deubiquitination. **C.** Diagrams displaying different chemical structure of the bond between UBE2N and Ub in UBE2N∼Ub and UBE2N-Ub. **D.** MavC-catalyzed transglutamination. The product UBE2N-Ub suppresses the activation of NF-κB. **E.** MavC-catalyzed deamidation of Ub in solution.

**Fig. S2.**
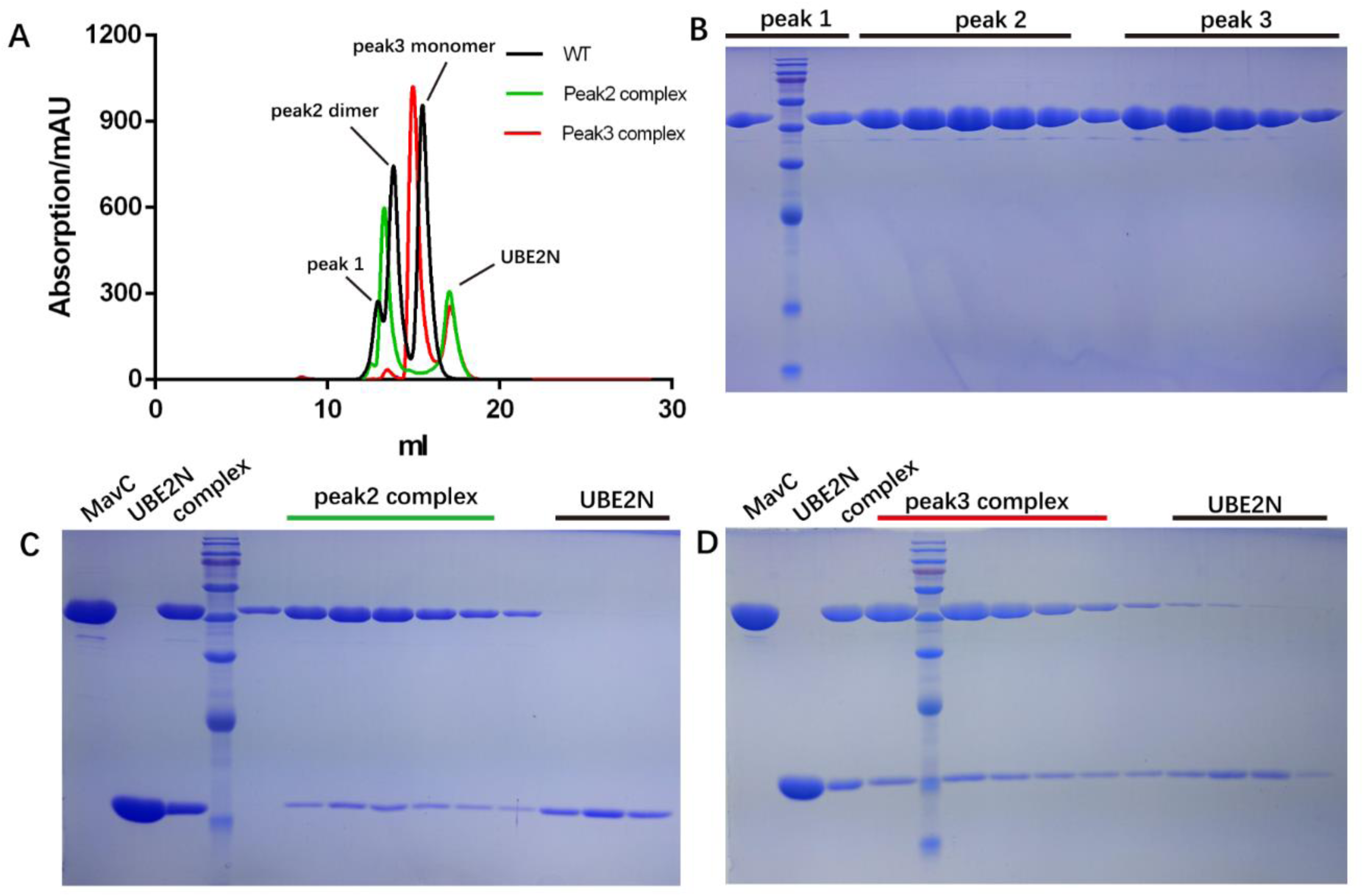
Two variants of UBE2N-Ub complexes used for crystallization. **A.** MavC eluted from peak 2 and peak 3 in gel filtration was incubated with UBE2N at a ratio of 1:2 and purified by size-exclusion chromatography using a Superdex200 increase column. The figure compares retention volumes of MavC (black), MavC (peak2) -UBE2N complex (green) and MavC (peak3) -UBE2N (red) complex. **B-D.** The elution peaks of MavC and MavC-UBE2N complexes were examined by SDS-PAGE. The word “complex” in the third lane 3 of gel images (C) and (D) is used to denote complex samples before gel filtration.

**Fig. S3.**
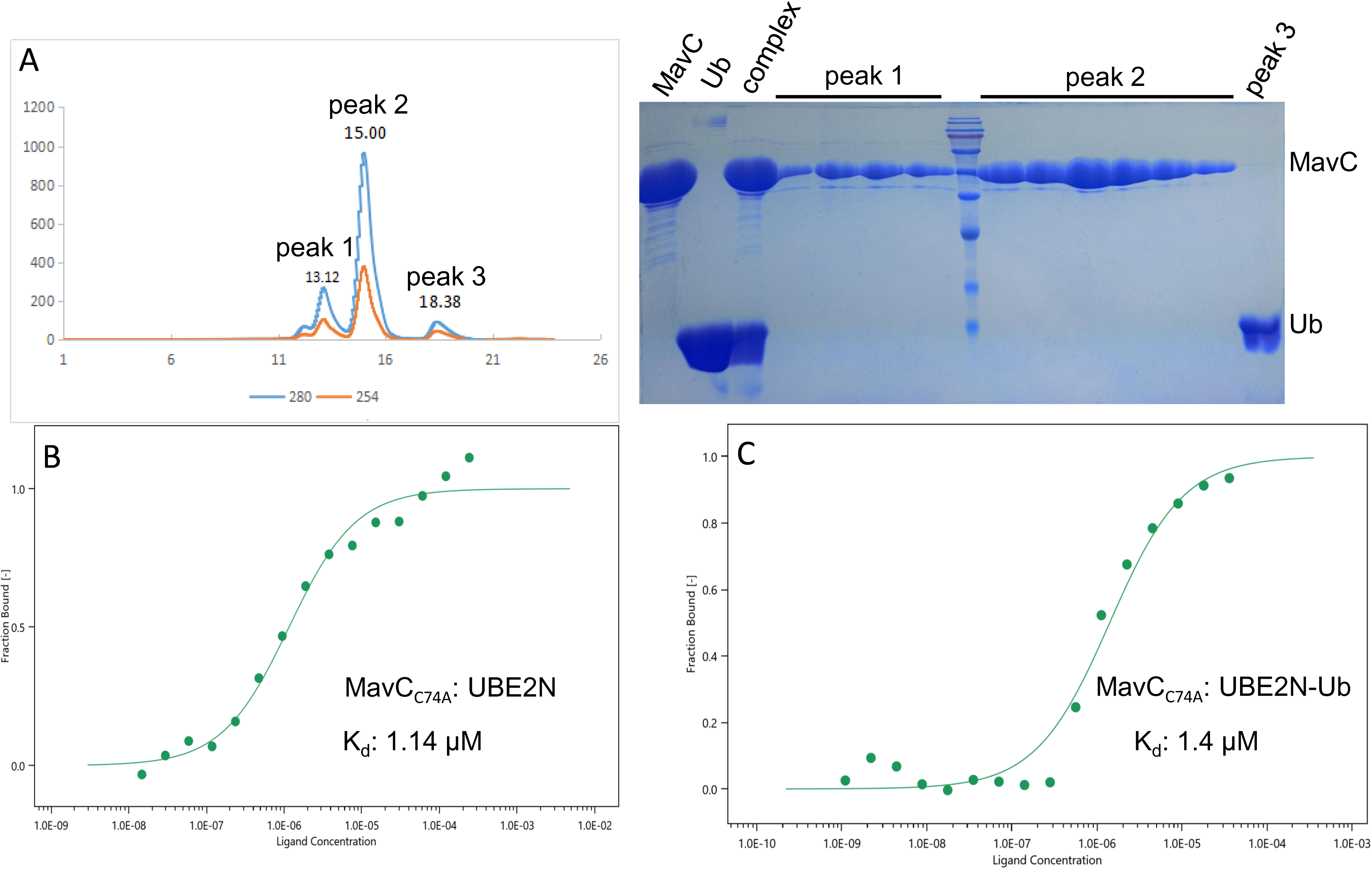
Interactions of MavC and Ub, MavC and UBE2N, and MavC and UBE2N-Ub in vitro. **A.** Purified MavC was incubated with Ub at a ratio of 2:1 and purified by size-exclusion chromatography using a Superdex200 Increase column. Peak 1 (dimeric MavC), peak 2 (monomeric MavC) and peak3 (Ub) were examined by SDS-PAGE. The word “complex” in the third lane 3 of gel images is used to denote complex samples before gel filtration. **B.** The interactions between MavC _C74A_ and UBE2N (panel B), and MavC _C74A_ and UBE2N-Ub (panel C) were measured by microscale thermophoresis (MST).

**Fig. S4.**
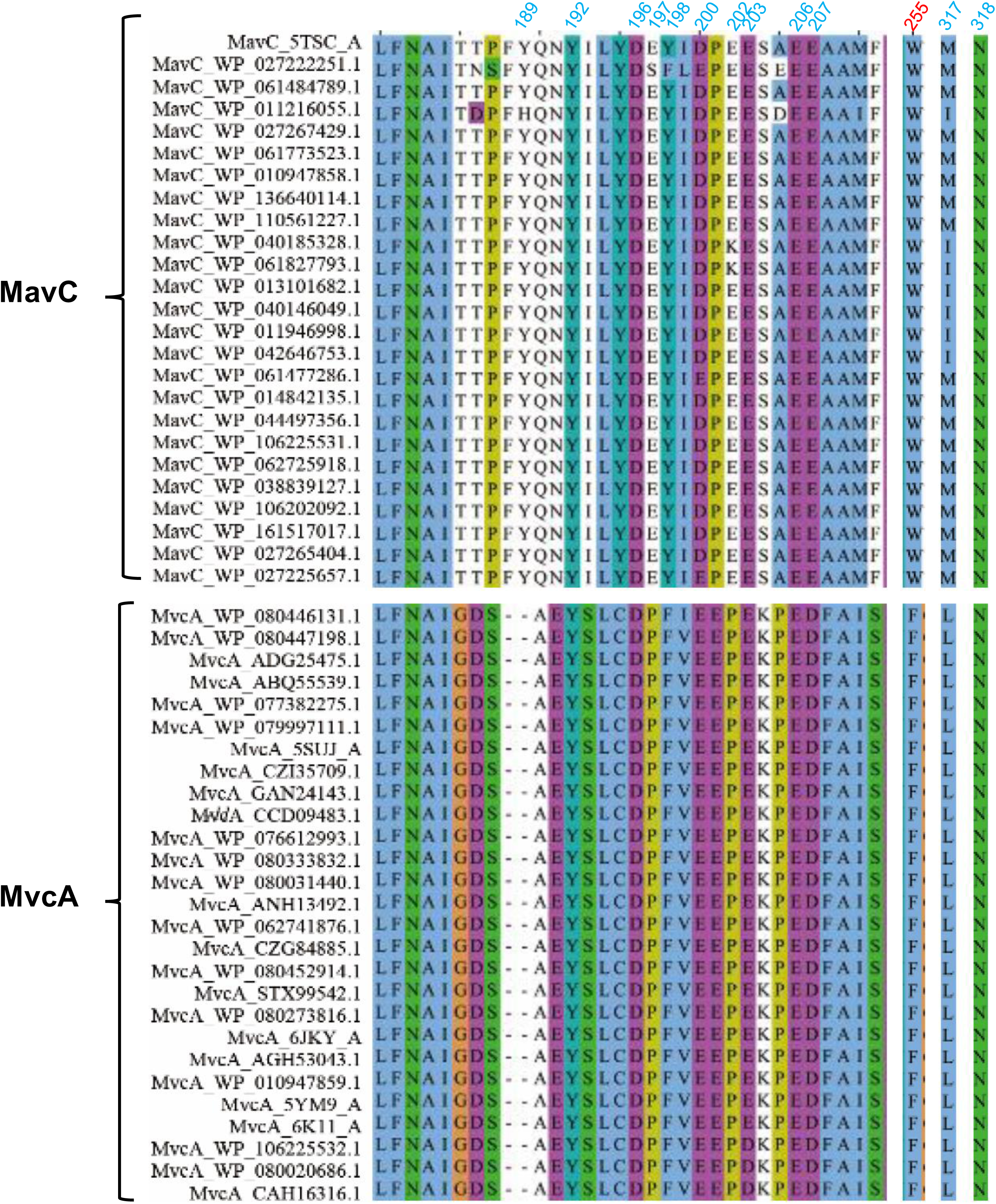
Multiple sequence alignment of MavC and MvcA sequences. MavC and MvcA sequences alignments carried out using MUSCLE v3.8.31 and visualized with Jalview v2.10.3. Residues involved in electrostatic interactions between the negative surface of MavC and positive surface of UBE2N are labeled with cyan. Trp255 of MavC in the active site is labeled with red.

**Fig. S5.**
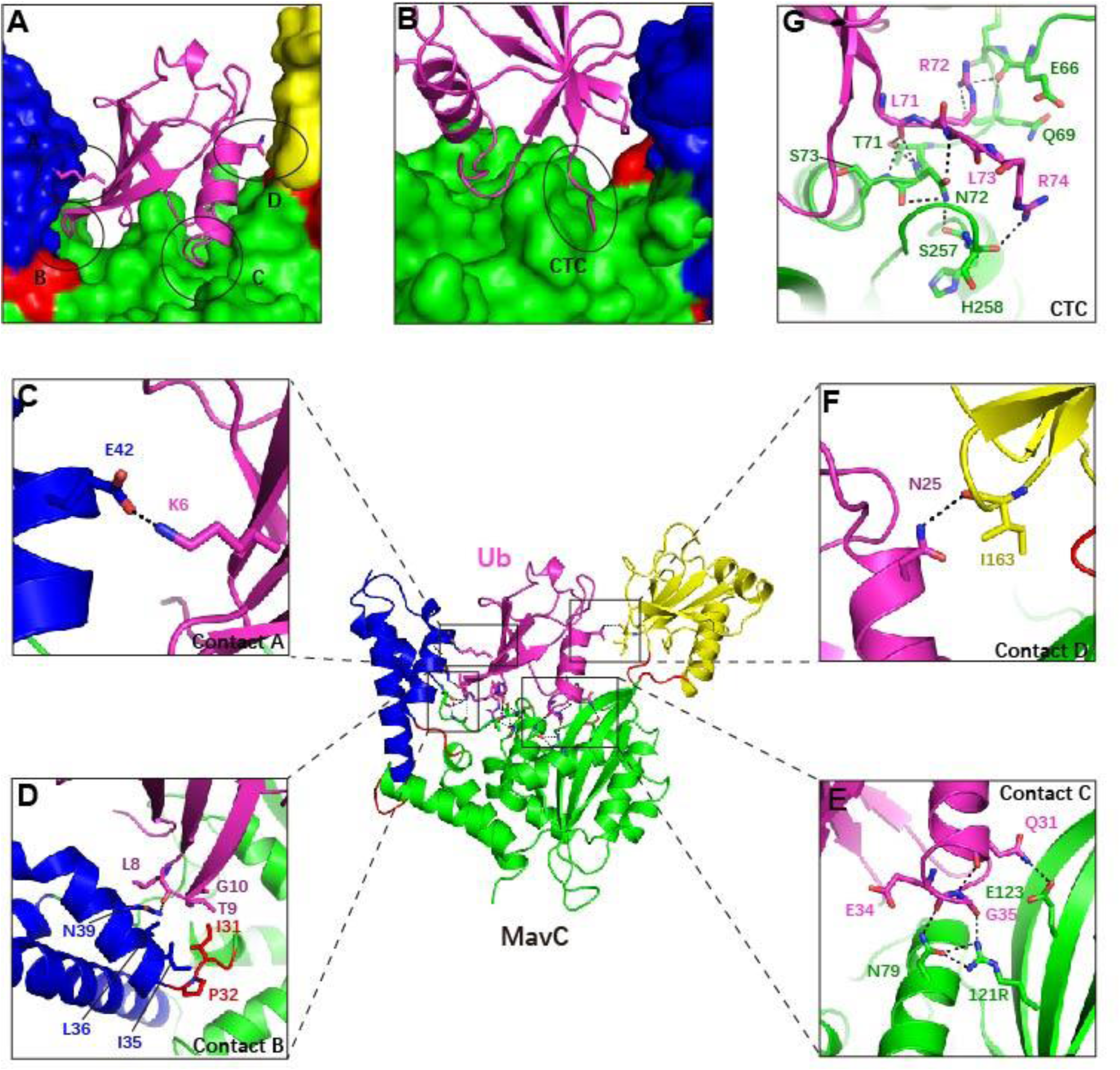
Interactions between MavC and Ub in the MavC-UBE2N-Ub complex. **A-B**. The five contacting interfaces between MavC (green-yellow-blue surface) and Ub (magenta cartoon) are indicated by circles. **C-G**. Detailed views of contact A (panel C), contact B (panel D), contact C (panel E), contact D (panel F), and the C-terminal contact (CTC) (panel G). Key residues involved in interactions between MavC (green-yellow-blue) and Ub (magenta cartoon) are shown as sticks. Hydrogen bonds are indicated by black dashed lines.

**Fig. S6.**
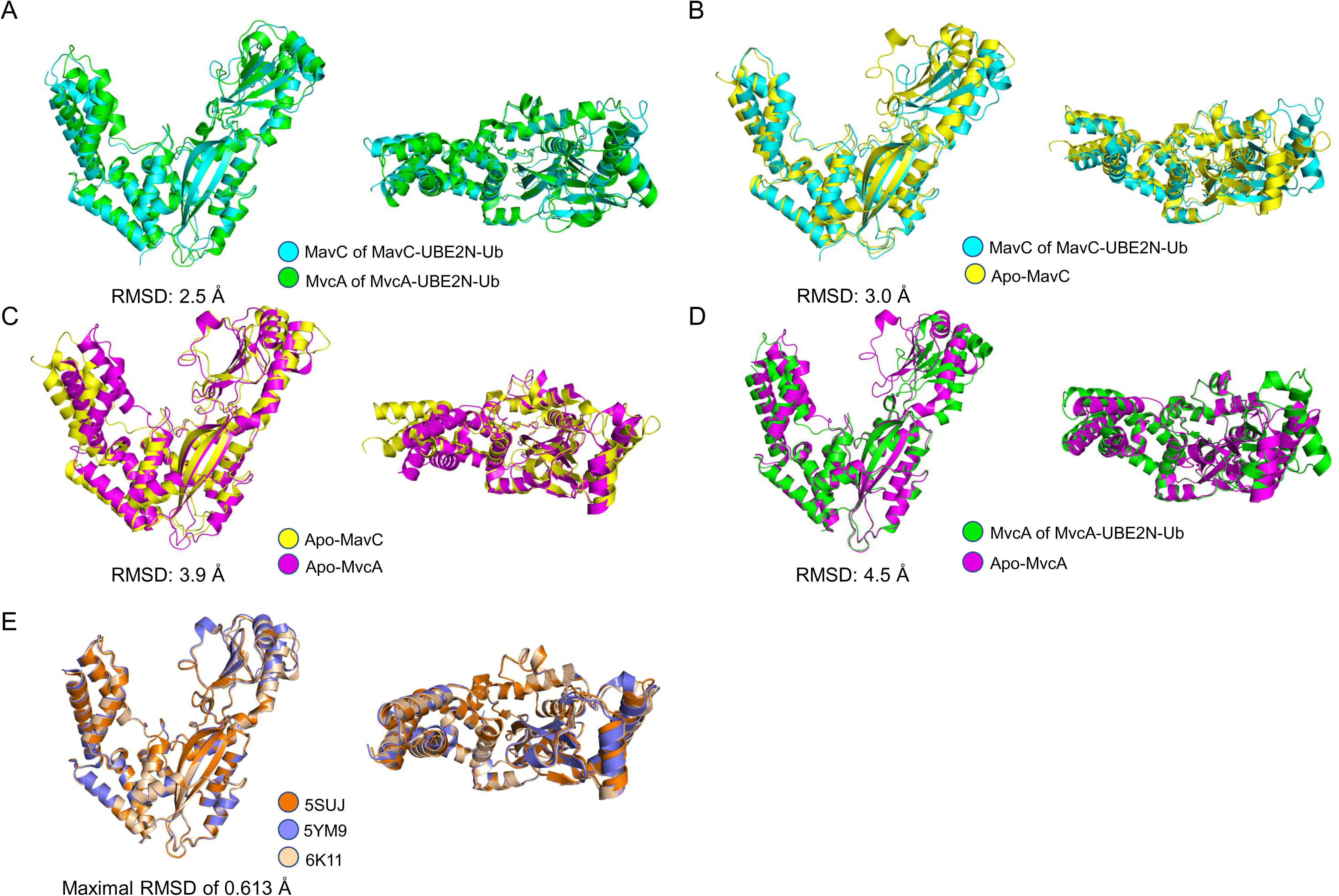
Structural alignments of MavC and MvcA in apo form and ternary complex. **A.** Structural alignment of the ternary complex of MavC-UBE2N-Ub (cyan) and MvcA-UBE2N-Ub (green) with an RMSD of 2.5 Å. **B.** Structural alignment of the MavC-UBE2N-Ub ternary complex (cyan) and apo MavC (yellow) with an RMSD of 3.0 Å. **C.** Structural alignment of apo MavC (yellow) and apo MvcA (magenta) with an RMSD of 3.9 Å. **D.** Structural alignment of the MvcA-UBE2N-Ub ternary complex (green) and apo MvcA (magenta) with an RMSD of 3.0 Å. **E.** Structural alignment of three available apo MvcA structures (PDB ID: 5SUJ, 5YM9, 6K11) with a maximal RMSD of 0.613 Å.

**Fig. S7.**
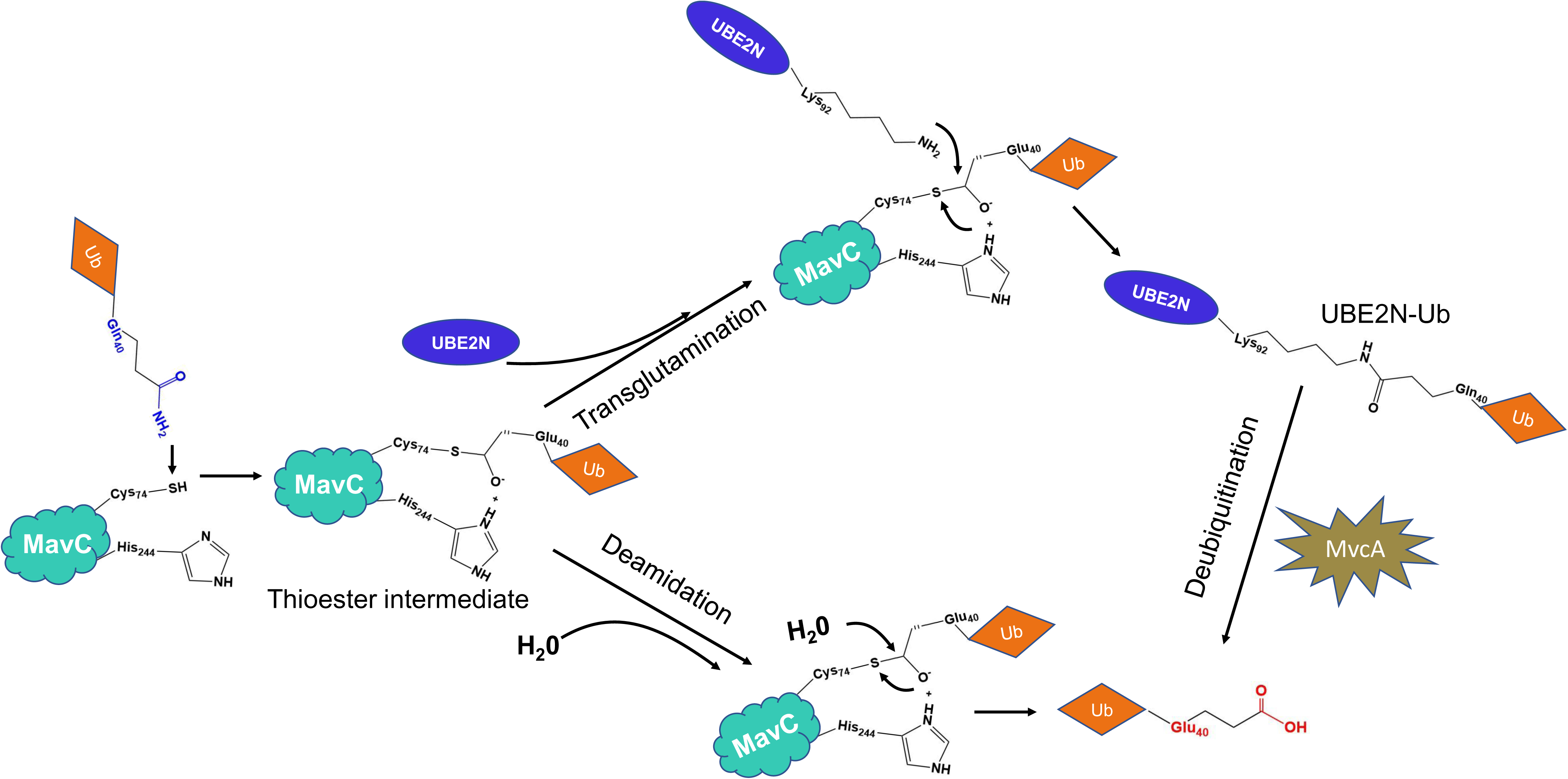
Proposed mechanism for transglutamination and deamidation reaction catalyzed by MavC. In the first step, Cys74 of MavC attacks Gln40 of Ub to generate a thioester intermediate. This intermediate reacts with Lys92 on UBE2N to form UBE2N-Ub or with H_2_O to form Glu40 in the absence of UBE2N. The ispoeptide bond in UBE2N-Ub complex catalyzed by MavC can be cleaved by MvcA.

## Methods

### Media, bacteria strains, plasmid constructions and cell lines

*Legionella* strains used in this papers were derivatives of *Philadelphia* 1 strain Lp02 and were grown and maintained on CYE (charcoal-yeast extract) plates or in AYE ^31^. For complementation experiments, the genes were cloned into pZL507 ^32^. *E.coli* strains XL1-Blue grown in Luria broth (LB) were used for expression and purification of all the recombinant proteins. Genes for protein purification were cloned into pQE30 (QIAGEN). Raw 264.7 and U937 cells were cultured in RPIM1640 medium in the presence of 10% FBS. When necessary, U937 cells were differentiated into macrophages by phorbol 12-myristate 13-acetate as described earlier ^33^. All cell lines were directly purchased from ATCC.

### Purification of proteins for biochemical experiments

For protein production, 10 mL overnight cultures were transferred to 200 mL LB medium in the presence of 100 µg ampicillin and grown to OD_600 nm_ of 0.6–0.8. The cultures were then incubated at 18°C for 16–18 h after adding isopropyl β-D-1-thiogalactopyranoside (IPTG) to a final concentration of 0.2 mM. Bacterial cells were harvested at 12,000x*g* by spinning and lysed by sonication. The soluble lysates were cleared by spinning at 12,000x*g* twice at 4°C for 20 min. His-tagged proteins were purified with Ni^2+^-NTA beads (Qiagen) and eluted with 300 mM imidazole in PBS buffer. Purified proteins were dialyzed with buffer containing 50 mM Tris-HCl (pH7.5), 150 mM NaCl, 5% glycerol, and 1 mM DTT.

### Purification of proteins for structural study

The gene of MavC (full length) was PCR amplified from *L. pneumophila* genomic DNA and inserted into pGEX-6p-1. The gene sequences of MavC_C74A_ (7-384 AA, Cys74 residue mutated to Ala) and UBE2N_K94A_ (full length) were also inserted into pGEX-6p-1. The gene sequence of Ub (1-76 AA) was inserted into pQE30. These plasmids were transformed into *E. coli* BL21(DE3) cells. The cells were grown in LB medium at 37°C with constant shaking at 220 rpm until the cell concentration reached OD_600_ of 0.8, after which the recombinant protein expression was induced by the addition of IPTG to a final concentration of 0.3 mM. The recombinant proteins were expressed at 18°C for 16 h. The cells were pelleted by centrifugation (5,000x*g*, 15 min) and subsequently resuspended in the cold lysis buffer (50 mM Tris-HCl pH 7.5, 150 mM NaCl). Following lysis by ultrasonication, the lysates were centrifuged at 17,000 rpm for 30 min at 4°C. The proteins with glutathione S-transferase (GST) tag or His tag were purified by affinity chromatography (glutathione agarose resin and Ni^2+^ resin, respectively). The GST tag was removed by the PreScission protease (PPase). The tag-less protein was then purified by size-exclusion chromatography (SEC) using a Superdex 200 increase column (GE Healthcare) equilibrated with a buffer containing 25 mM HEPES, pH 7.5, 150 mM NaCl and 2 mM DTT.

To prepare the MavC_C74A_-UBE2N_K94A_ complex, UBE2N_K94A_ was incubated with MavC_C74A_ at 2:1 molar ratio at 4°C for 1 h. To cross-link UBE2N_K94A_ and Ub, wild-type MavC was incubated with UBE2N_K94A_ and Ub at a molar ratio of 1:120:180 in a reaction buffer containing 25 mM HEPES, pH 7.5, 150 mM NaCl, 2 mM DTT and 10 mM Mg^2+^ at 25°C for 5 min. The UBE2N_K94A_-Ub was separated from MavC and excess UBE2N by Ni^2+^-NTA beads at 4°C and then purified by gel filtration to remove the excess Ub ^21^. The UBE2N_K94A_-Ub was collected and incubated with MavC_C74A_ at 1:2 molar ratio at 4°C for 1 h. Finally, the MavC_C74A_-UBE2N-Ub complex was separated by gel filtration, pooled and concentrated to A_280_=14 for the use in crystallization screen.

### Crystallization, data collection and structural determination

Crystallization screens were performed using the sitting drop vapor diffusion method at 16°C, with drops containing 0.5 µl of the protein solution mixed with 0.5 µl of reservoir solution. Diffraction-quality MavC_C74A_-UBE2N_K94A_-Ub crystals were obtained in 0.2 M sodium malonate pH 6.0, 20% w/v PEG 3,350, whereas MavC_C74A_-UBE2N_K94A_ crystals were obtained in 0.2 M Sodium malonate pH 7.0, 20% w/v PEG 3,350. The crystals were harvested and flash-frozen in liquid nitrogen with 20% glycerol as cryoprotectant. Complete X-ray diffraction datasets were collected at BL-17U1 beamline of Shanghai Synchrotron Radiation Facility (SSRF). Diffraction images were processed with the HKL-2000 program^34^. Structures of the binary and ternary complex were solved by molecular replacement (MR) using Phaser-MR (MavC: 5TSC, UBE2N: 1JAT, Ub: 4HCN) ^23, 35–37^. Model building and crystallographic refinement were carried out in Coot and PHENIX ^38, 39^. The interactions were analyzed with PyMol (http://www.pymol.org/) and PDBsum. The figures were generated in PyMol. Detailed data collection and refinement statistics are listed in Table 1.

### MavC-induced ubiquitination and deamidation assays

For MavC-mediated ubiquitination of UBE2N, 5 µg Ub, 0.5 µg UBE2N or its mutant derivatives and 0.05 µg MavC or its mutant derivatives were incubated for 8 min or 30 min at 37°C in 25 µl reactions systems containing 50 mM Tris-HCl (pH7.5), 5 mM Mn^2+^ and 1 mM DTT. Reaction products were resolved by SDS-PAGE, protein bands were visualized by Coomassie brilliant blue staining and the amount of protein in bands was quantified by densitometry.

For MavC-mediated deamidation of Ub, 10 µg Ub and 1 µg MavC or its mutant derivatives were incubated for 1 h at 37°C in 25 µl reactions containing 50 mM Tris-HCl (pH8.8). Reaction mixtures were then mixed with 5x native gel loading buffer and resolved by native PAGE followed by Coomassie brilliant blue staining.

### MavC-mediated ubiquitination of UBE2N during *L. pneumophila* infection

For *L. pneumophila* infection experiments, all *Legionella* strains including complementation strains were grown overnight in AYE medium to post-exponential phase (OD_600nm_=3.2-3.8) and were induced with 0.2 mM IPTG for 3 h at 37°C before infection. Raw264.7 or U937 cells were infected with *L. pneumophila* strains at a MOI of 10 for 2 h. Cells washed with PBS three times were collected and lysed with 0.2% saponin on ice for 30 min. The cell lysates were resolved by SDS-PAGE and probed with MavC-specific antibody to check the translocation of MavC and its mutants, and modified UBE2N was probed with UBE2N-specific antibody.

